# Immunogenetic metabolomics revealed key enzymes that modulate CAR-T metabolism and function

**DOI:** 10.1101/2023.03.14.532663

**Authors:** Paul Renauer, Jonathan J. Park, Meizhu Bai, Arianny Acosta, Won-Ho Lee, Guang Han Lin, Yueqi Zhang, Xiaoyun Dai, Guangchuan Wang, Youssef Errami, Terence Wu, Paul Clark, Lupeng Ye, Quanjun Yang, Sidi Chen

## Abstract

Immune evasion is a critical step of cancer progression that remains a major obstacle for current T cell-based immunotherapies. Hence, we seek to genetically reprogram T cells to exploit a common tumor-intrinsic evasion mechanism, whereby cancer cells suppress T cell function by generating a metabolically unfavorable tumor microenvironment (TME). Specifically, we use an *in silico* screen to identify *ADA* and *PDK1* as metabolic regulators, in which gene overexpression (OE) enhances the cytolysis of CD19-specific CD8 CAR-T cells against cognate leukemia cells, and conversely, *ADA* or *PDK1* deficiency dampens such effect. *ADA*-OE in CAR-T cells improves cancer cytolysis under high concentrations of adenosine, the ADA substrate and an immunosuppressive metabolite in the TME. High-throughput transcriptomics and metabolomics in these CAR-Ts reveal alterations of global gene expression and metabolic signatures in both *ADA-* and *PDK1-* engineered CAR-T cells. Functional and immunological analyses demonstrate that *ADA*-OE increases proliferation and decreases exhaustion in α-CD19 and α-HER2 CAR-T cells. ADA-OE improves tumor infiltration and clearance by α-HER2 CAR-T cells in an *in vivo* colorectal cancer model. Collectively, these data unveil systematic knowledge of metabolic reprogramming directly in CAR-T cells, and reveal potential targets for improving CAR-T based cell therapy.

**Synopsis:** The authors identify the adenosine deaminase gene (ADA) as a regulatory gene that reprograms T cell metabolism. ADA-overexpression (OE) in α-CD19 and α-HER2 CAR-T cells increases proliferation, cytotoxicity, memory, and decreases exhaustion, and ADA-OE α-HER2 CAR-T cells have enhanced clearance of HT29 human colorectal cancer tumors *in vivo*.

## Introduction

Cancer cells have several mechanisms to suppress immune function, including the development, recruitment, and activity modulation of suppressive immune cells. In addition, the tumor microenvironment (TME) itself can suppress T cell function through the oxygen depletion (hypoxia), nutrient deprivation, and production of immunosuppressive metabolites, including lactic acid and adenosine [1,2]. Cancer cells can adapt to these unfavorable conditions through metabolic reprogramming, offering competitive advantage over tumor-infiltrating T cells. Metabolic fitness is also a critical determinant of CTL cytotoxicity, persistence, and exhaustion [3], which remain challenges for improving anti-cancer efforts of CTLs within the TME. In addition, there is significant knowledge gap between the metabolism of T cells and chimeric antigen receptor (CAR) T cells, patient-derived T cells that are reprogrammed to target specific cancer antigens and reintroduced to patients as immunotherapy [4,5]. The metabolism of these cells may differ because of the highly engineered chimeric antigen receptor and the rewired T cell signaling that is used by the receptor [6].

Therefore, we hypothesize that we can improve CAR-T cell function through genetic manipulation of key metabolism genes in T cells. In this study, we screened genes that potentially influence T cell metabolism in the TME by a rational selection process through large databases, literature, and metabolic pathway analyses to narrow down to a small number of candidates. Through gene editing, systems biology approaches, and metabolomics experiments, (1) we revealed and validated key enzymes that modulate T cell and CAR-T metabolism; (2) we characterized the transcriptomic, metabolomic, and functional changes to T cell and CAR-T cell phenotypes; and (3) explored the anti-cancer efficacy of the reprogrammed CAR-T cells, which demonstrated enhanced cytotoxic functions, increased memory, and decreased exhaustion.

## Materials and Methods

### Institutional Approval

This study has received institutional regulatory approval. All recombinant DNA and biosafety work were performed under the guidelines of Yale Environment, Health and Safety (EHS) Committee with an approved protocol (Chen-rDNA-15-45; Chen-rDNA-18-45). All human sample work was performed under the guidelines of Yale University Institutional Review Board (IRB) with an approved protocol (HIC#2000020784). All animal work was performed under the guidelines of Yale University Institutional Animal Care and Use Committee (IACUC) with approved protocols (Chen 2018-20068; 2021-20068).

### Murine tumor models

NOD-SCID IL2Rγ-null (NSG) mice were purchased from JAX and bred for *in vivo* tumor models. Briefly, 2e6 HT29-GFP-Luciferase (HT29-GL) cells were subcutaneously injected into the right flanks of mice, and tumors were measured by caliper with volume = (0.5 x height x width x depth). After 10 days, tumors reached an average volume > 75 mm3, and were treated with 3e6 CD3+ T cells, α-HER2 CAR-T, or ADA-OE α-HER2 CAR-T cells via tail-vein injection (2 independent experiments of 4-5 mice per condition group). Tumor volume measurements were then recorded every 2-3 days for the duration of the experiments, after which, the tumors were extracted into PBS supplemented with 2 % FBS. The tissue was macerated, incubated with collagenase IV for 1 hour at 37C, then transferred through a 40 μm filter prior to staining the cells for flow cytometry analysis.

### Selection of metabolic regulatory genes

Metabolic regulatory genes were predicted using a multi-stage selection process starting with genes that might have a potential role in T cell functions, which were pooled from three different gene-set catalogs, including genes associated with tumor-immunity, autoimmunity, and human immune-related diseases (ImmunoBase; https://genetics.opentargets.org/immunobase; accessed: 11/2018). The pooled genes were filtered for (1) an association with T cell metabolism, (2) enzymes that interacted with TME-related metabolites, (3) enzymes with easily detected substrates based on published data, (4) genes that were disrupted by CRISPR-RNP with any of three top gRNAs designed by the GPP web portal [7,8], and (5) enzymes with detectable substrate changes (LC/MS) in HEK293FT treated with CRISPR-RNP gRNA knockout vs NTCs. All gene lists and references used for the candidate gene selection process are detailed in **Supplementary Data 1**.

### Regular cell culture

HEK293FT cell lines were obtained from ThermoFisher and cultured in D10 media (DMEM (Gibco) supplemented with 10 % FBS (Sigma-Aldrich) and 200 U / mL penicillin–streptomycin (Gibco)). NALM6 were obtained from American Type Culture Collection (ATCC) and cultured in cRPMI media (RPMI-1640 (Gibco) supplemented with 10 % FBS (Sigma-Aldrich), and 200 U / mL penicillin-streptomycin (Gibco). NALM6-GL (GFP and luciferase reporters) were produced via lentiviral infection using a GFP-luciferase reporter vector, as previous described [9].

### Cytotoxicity assays

The co-culture cytotoxicity assays were performed by seeding 96-well flat-bottom culture plates with NALM6-GL cells at 2e4 cells / well, and T cells were added at different concentrations for 48 h in X-VIVO^TM^ 15 media (Lonza) supplemented with 5 % human AB serum (Gemini Bio) and 30 U / mL recombinant human IL-2 (BioLegend). The viability of NALM6-GL cells was then assessed by adding 150 ug / mL D-Luciferin (PerkinElmer) into each well for 10 mins at 37 °C, then measuring the relative luciferase intensity via plate reader (PerkinElmer).

### CD8 T cell isolation and culture

Human primary peripheral blood samples were derived from healthy donors (STEMCELL). CD8 T cells were purified using the EasySep human CD8 T cell isolation kits (STEMCELL), as per manufacturer’s instructions (STEMCELL). Cell culturing was performed in X-VIVO^TM^ 15 media (Lonza), supplemented with 5 % human AB serum (Gemini Bio) and 30 U / mL recombinant human IL-2 (BioLegend) [9], and cells were activated for 16 h with anti-human CD3 / CD28 Dynabeads at 1:1 ratio, using manufacturer’s instructions (Gibco). Subsequently, activated T cells were cultured for another 48 h before ribonucleoprotein (RNP) electroporation or lentiviral infection, described below.

### Gene editing via Cas9-crRNA:tracrRNA RNP electroporation

All guide RNAs (gRNAs) were designed using the Broad Genetic Purtubation Platform Web Portal [7,8] and tracrRNA and crRNA were synthesized by Integrated DNA Technologies, Inc. For genetic editing, human CD8 T cells were first activated with human anti-CD3 / CD28 Dynabeads as above. The T cells were transfected using the Neon Transfection System (Invitrogen) and Neon Transfection System Kit (Thermo-Fisher), as per manufacturer’s instructions. Briefly, gRNAs were made by annealing crRNA and tracrRNA at a 1:1 ratio in a thermocycler, using the following settings: 95 °C for 5 min, slowly cool down at -5 °C / min, then hold at 25 °C. Next, RNP were formed by incubating the gRNA and HiFi Cas9 Protein v3 for 15 min at room temperature, then the RNP mixed with electroporation enhancer (Final concentration: 1.8 uM gRNA, 1.5 uM Cas9) and 2e6 T cells / 100 uL. The mixture was electroporated using the Neon Transfection System with the setting of 1,600 V and 10 ms for 3 pulses, then subsequently transferred T cells to pre-warmed supplemented X-VIVO^TM^ 15 media for continued culturing.

### Vector design

The anti-CD19 chimeric antigen receptor (CAR) construct is comprised of a leader sequence, a FLAG-tag sequence (GATTACAAAGACGATGACGATAAG), a human antigen-binding domain of the FMC-63 single chain fragment variable (scFv) that targets CD19 (GenBank: HM852952), and a CD8 hinge and transmembrane domain linked to the 4-1BB (CD137) transactivation domain and CD3 zeta signaling domain (**Fig. 3a**) [9,10]. The CAR constructs sequences were cloned into lentiviral plasmid backbone pLY041a (EF1α-T2A-GFP-WPRE) where T2A-GFP was replaced with the CAR sequence. Additional CAR constructs were created for the tandem expression of either *ADA* or *PDK1* cDNA, for which genes were cloned from commercial cDNA plasmids (MHS6278-202829515 for *ADA* and OHS1770-202322779 for *PDK1*; Horizon Discovery Ltd) into a multi-cistronic vector, downstream of the CAR and a T2A self-cleaving peptide sequence via the Gibson Assembly method [11].

### Lentivirus production

Lentivirus particles were produced from HEK293FT cells, as previously described [12]. Briefly, the HEK293FT cells were transfected in 15 cm culture dishes with a mixture of 20 μg target plasmid, 15 μg psPAX2 (Addgene) packaging plasmid, 10 μg pMD2.G (Addgene) envelope plasmid and 130 μL 1 μg / mL polyethylenimine (Sigma-Aldrich) in 450 μL Opti-MEM media (Lonza). The virus was collected at 48 h and 72 h post-transfection, then concentrated with AmiconUltra-100kD Ultrafiltration centrifuge tubes (Millipore). CD8 T cells were spin-infected with lentivirus at 900 x *g*, 32 °C for 60 min in 24-well tissue-culture plates, with each well containing 200 μL lentivirus and 800 μL supplemented X-VIVO^TM^ 15 media including 2.4e6 cells and polybrene. Following spin-infection, media was completely replaced 16 h later, and cells were maintained at a density of 1-2e6 cells / mL.

### Measurement of gene editing

Genetic perturbations from CRISPR-RNP were quantified from genomic DNA (gDNA) that was extracted from T cells using QuickExtract Buffer (NEB) according to manufacturer’s recommendation. The genomic regions targeted by CRISPR-RNP were amplified by PCR (95 °C / 3 min; 36 cycles: 95 °C / 30 s, 60 °C / 30 s and 72 °C / 45 s; 72 °C / 4 min), 1 μL QuickExtract Buffer digested mixture was used as template in a 50 μL PCR reaction with DreamTaq Green 2X PCR Master Mix (Thermo Fisher Scientific) and 0.5 μM oligos. For T7 endonuclease I (T7eI) assay [13], 200 ng of purified PCR product was denatured at 95 °C for 5 min, then reannealed to form heteroduplexes (cooled to 25 °C at -5 °C / min), digested annealed DNA with 10 U of T7eI (NEB) at 37 °C for 45 min, then visualized on a 2 % agarose E-Gel with SYBR-Gold (Invitrogen). DNA fragment sizes were quantified using an ImageJ software [14], and cutting efficiencies were calculated as the proportion of T7eI-digested bands. For Nextera sequencing, 0.5 ng of PCR product was used for tagmentation and subsequently barcoded, as per manufacturer’s instructions (Illumina). The processed DNA was pooled, purified by 2 % agarose E-Gel electrophoresis, extracted from the gel via DNA Gel Extraction kit (Qiagen), and the pooled DNA was sequenced by the Illumina Novaseq platform (100 bp paired-end; Illumina). For indel quantification, paired reads were mapped to the amplicon sequences using BWA-MEM with default settings (BWA-v0.7.1) [15,16], sorted and indexed using Samtools [17], variants were called using mpileup via Samtools with a max depth of 100000 [18,19], and indels were quantified using Varscan pileup2indel command [20] and filtered (>= 2 indel reads, >=2 coverage at indel locus, indel frequency > 0.00001). Cutting efficiency was reported as the proportion of indels detected with a 75 bp window of the target cut site (**Supplementary Data 2**).

### Quantitative RT-PCR and mRNA-seq

Total RNA was extracted by treating cells with TRIzol Reagent (Invitrogen), then isolating RNA with the Qiagen RNA extraction kit, as per manufacturer’s instructions. For the qRT-PCR, the SuperScript™ IV Reverse Transcriptase (Invitrogen) was used for cDNA synthesis, and target mRNA levels were quantified relative to ACTB via qPCR with Taqman probes (Invitrogen) (**Supplementary Data 2**). For mRNA-seq, library preparation was performed using the NEBNext® Poly(A) mRNA Magnetic Isolation module (Illumina) and NEBNext® Ultra™ II Directional RNA Library Prep Kit (Illumina), per manufacturer’s instruction. Briefly, all samples were normalized to 200 or 500 ng total RNA for fragmentation and cDNA synthesis. Double-stranded cDNA was purified using oligo-dT Sample Purification Beads, adapters were ligated to the cDNA, and samples were barcoded using the NEBNext® Multiplex Oligos for Illumina® (Index Primers Set 1) (NEB), listed in the **Supplementary Data 1**. Sample barcoding PCR conditions were 1 cycle of 98 °C for 30 s, 11 cycles of 98 °C for 10 s, 65 °C for 75 s, and one cycle of 65 °C for 5 min. The barcoded library was then purified using NEBNext Sample Purification Beads then sequenced using the Novaseq platform (Illumina).

### Protein extraction and Western Blot

Cellular protein was extracted by treating 1e7 cells / mL in ice-cold 1 x RIPA buffer (Boston BioProducts), containing protease inhibitors (Roche, Sigma) for 30 min on ice, pelleting cell debris for 15 min at 12,000 x *g* at 4 °C, then the protein-containing supernatant was collected, and concentrations were measured by BCA assay (Abcam). Proteins were denatured in Laemmli loading buffer at 95 °C for 5 min, and 20 μg / sample were loaded into 4 % – 20 % SDS-PAGE gel for electrophoresis at 80 V for 20 min then 120 V for 70 min. Proteins were transferred to PVDF membranes using a wet transfer at 320 mA for 70 min on ice. Membranes were blocked at room temperature for 1 h using 3 % BSA in TBST buffer, transferred membranes were incubated in 1:1000 diluted primary antibody in TBST buffer with 3 % BSA at 4 °C overnight, then incubated with 1:5000 horseradish peroxidase (HRP)-conjugated secondary antibodies in TBST buffer for 1 hr at room-temperature. The chemiluminescent substrate (Bio-Rad) was added to membranes and immunoblots were imaged using an Amersham Imager 600 (GE Healthcare).

### mRNA-seq data processing

mRNA sequencing data was aligned to the human transcriptome reference panel (GRCh38 Ensembl Version 96) and quantified from raw FASTQ files using Kallisto with the quant function and 100 bootstraps [21]. Differential expression analyses were performed using the Sleuth package for R [22], by fitting the data, first to a full measurement error model then a secondary reduced error model, and lastly, data were analyzed for differential expression and beta-values via the Wald test. Differentially expressed genes (q value < 0.05 and beta-value fold-change > 1) were analyzed by the Database for Annotation, Visualization and Integrated Discovery (DAVID) gene set enrichment test [23,24], and results were filtered to include Biological Process Gene Ontology terms, and Benjamini-adjusted p values < 0.05. Quantitative enrichment analyses were performed by the fast Gene Set Enrichment Analysis (GSEA) package for R, using the beta-values of differentially expressed genes (q < 0.01) and pathways from the MSigDB hallmark collection (MSigDB v7.0) [25,26] with 1000 permutations, and results were filtered to include those with a gene set size between 10 and 200. Results were visualized using the ggplot2 R package.

### Flow cytometry

Cell surface proteins were stained with experimentally-titrated fluorophore-conjugated antibodies in MACS buffer (2 % BSA and 2 mM EDTA in PBS) for 30 min on ice. Stained cells were washed then resuspended in MACS buffer for flow cytometric analysis. For intracellular staining, cells were fixed for 20 min on ice with fixation / permeabilization buffer (Invitrogen), washed twice with permeabilization buffer (Invitrogen), stained with experimentally-titrated fluorophore-conjugated antibodies in permeabilization buffer (Invitrogen) for 30 min on ice, then washed and resuspended in MACS buffer. Cytometric analyses were performed using an Attune NxT Focusing Flow Cytometer (Thermo Fisher Scientific), and cell sorting was accomplished with a FACSAria II flow sorter (BD) and cytometry data was analyzed using FlowJo software v10.3 (Treestar, Ashland, OR).

### Metabolomics

Metabolomics analyses were performed using 1e6 CD3^+^CD8^+^ T cells that were purified via FACS and washed twice with DPBS. Lysates were extracted by treating the cells with 800 μL of 80 % (vol / vol) methanol (precooled to -80 °C) on dry ice for 30 min then ice for 20 min, and lysate was separated from cellular debris by centrifugation at 15,000 x *g* for 15 min at 4 °C twice to isolate the supernatant [27]. The lysate / methanol mixture was dried by Speedvac, resuspended in 80 % (vol / vol) methanol, centrifuged at 18,000 x *g* for 10 min to remove particulates, and stored at -80 °C. Metabolites were analyzed using the Agilent 6550 Q-TOF LC / MS system, then the Agilent 6490 Triple quadrupole LC / MS System. The HILIC liquid chromatography was optimized with the bioZen™ 2.6 µm Glycan LC Column, 150 x 2.1 mm (Phenomenex) and a Glycan guard column, 4 x 2 mm (Phenomenex). The eluents were A: 0.01 % formic acid in water HPLC grade and B: 0.01 % formic acid in acetonitrile. The gradient was set as follows: 0 – 2 min 94 % B; 2-8 min 94 – 90 % B; 8 – 16 min 90 – 76% B; 16 – 36 min, 76 – 50 % B; 36 – 42 min hold at 50 % B and then returned to initial conditions for 2 min for column equilibration. The flow rate was set at 0.3 mL / min.

### Metabolomics data analyses

Metabolomics quantification was performed using a two-part strategy, including an initial untargeted analysis to identify detectable metabolites across samples, and subsequent targeted analyses that re-evaluated these metabolites using standards. Briefly, our untargeted metabolomics analysis had an optimized workflow, consisting of automated peak detection and integration, peak alignment, background noise subtraction, and multivariate data analysis for comprehensive metabolite phenotyping of each experimental group using Agilent Mass Hunter Qualitative Analysis Software (Version B 6.0.633.0) and Agilent Mass Profiler Professional (Version B 6.0.633.0). The untargeted metabolites were identified based on accurate mass match (accurate mass error ±30 ppm) and fragmentation pattern match, while structural annotations were matched to the metabolite databases: HMDB (http://www.hmdb.ca/) and METLIN (http:// metlin.scripps.edu) using the mass-to-charge ratios. We established detectable metabolites as those that significantly differed in the gRNA T cells relative to the NTC T cells (q < 0.05, log2-FC > 0.5; comparative analysis is described in more detail below). The targeted metabolomics analyses were performed for our list of detectable metabolites, as well as metabolites from related proline metabolism and immune system metabolism pathways. The targeted metabolite quantification was performed as above with Multiple Reaction Monitoring (MRM) used for qualitative and quantitative analysis of purified standards (Sigma).

All metabolomics analyses were performed via MetaboAnalyst v4.0 [28–30], whereby raw peak intensities were filtered to remove low-detection features (> 50% missing values across samples), imputed to replace missing values (half of minimum value across samples), then filtered to remove near-constant values (5% of features with lowest interquantile range). Next, the data were sample-normalized to the pooled control group, log2-transformed and scaled to z-score. Differential metabolite levels were then calculated by unpaired two-tailed *t* test, assuming equal variance between groups. Metabolomics Pathway Analyses (MetPA) were performed with default settings: global enrichment test with pathway topologies calculated as relative-betweeness centrality, and the KEGG Homo sapiens pathway library (Oct 2019). Integrated transcriptome-metabolome MetPA were performed with log2-fold-changes (LFC) of significantly changed metabolites (q < 0.10) and the beta-value of differentially expressed genes (q < 0.01, abs(b) > 1) as input, using hypergeometric tests for enrichment with KEGG Homo sapiens metabolic pathways (Oct 2019), pathway topologies calculated as degree centrality, and unweighted integration of gene-metabolite p values. Significantly enriched pathways were those with an FDR-corrected p value < 0.05, and a pathway impact score > 0.

### Sample size determination

Sample size was determined according to the lab’s prior work or from published studies of similar scope within the appropriate fields.

### Replication

Number of biological replicates (usually n >= 3) are indicated in the figure legends. Key findings (non-NGS) were replicated in at least two independent experiments. NGS experiments were performed with biological replications as indicated in the manuscript.

### Randomization and blinding statements

*In vivo* experiments were randomized and blinded from the experimenter that measured tumor volumes. Regular *in vitro* experiments were not randomized or blinded. NGS data were processed in an unbiased manner using barcoding and metadata.

### Standard statistical analysis

Standard statistical analyses were performed using regular statistical methods. GraphPad Prism, Excel and R were used for all analyses. Different levels of statistical significance were accessed based on specific p values and type I error cutoffs (0.05, 0.01, 0.001, 0.0001). Further details of statistical tests were provided in figure legends and/or supplemental information.

### Data Collection summary

Flow cytometry data was collected by BD FACSAria or ThermoFisher Attune.

All deep sequencing data were collected using Illumina Sequencers at Yale Center for Genome Analysis (YCGA).

Co-culture killing assay data were collected with PE Envision Plate Reader.

### Data analysis summary

Flow cytometry data were analyzed by FlowJo v.10.

All simple statistical analyses were done with Prism 9.

All NGS analyses were performed using custom codes.

### Code availability

The code used for data analysis and the generation of figures related to this study are available from the corresponding author upon reasonable request.

### Data and resource availability

All data generated or analyzed during this study are included in this article and its Supplementary Data files. Specifically, data and statistics for non-high-throughput experiments such as flow cytometry, qPCR, protein experiments, and other molecular or cellular assays are provided in Supplementary Data Tables. Processed data for genomic sequencing (e.g. RNA-seq, amplicon sequencing) and other forms of high-throughput experiments (e.g. metabolomics) are provided as processed quantifications in the Supplementary Data Tables. Raw sequencing data have been deposited to the NIH Sequence Read Archive (SRA) and Gene Expression Omnibus under the series accession: GSE225186.

Review information:

Link: https://www.ncbi.nlm.nih.gov/geo/query/acc.cgi?acc=GSE225186

Accession: GSE225186

Token: yvyraomctfkhlqx

Original cell lines are available at commercial sources listed in supplementary information files. Genetically modified cell lines are available via Chen lab. Other relevant information or data are available from the corresponding author upon reasonable request.

## Results

### Candidate metabolic regulators of CD8+ T cells identified from human disease database and metabolic pathway analyses

To identify a set of promising candidate genes as genetic targets to metabolically reprogram T cells, we developed a multi-step rationalization and selection process (**Fig. 1a**). Briefly, a candidate list was constructed from 572 genes with potential roles in T cell function, chosen from the data-mining of tumor immunity studies (137 genes), the ImmunoBase catalog of genes relevant to human immune-related diseases (311 genes; https://genetics.opentargets.org/immunobase, ImmunoBase accessed: 11/2018), and genes that were associated with autoimmune diseases in genome-wide association studies (GWAS; 196 genes; **Fig. 1A; Supplementary Data 1; Methods**). The candidate gene list was then sequentially filtered, first for gene products that participate in metabolic pathways that are known to be related to T cell function (resulted in 67 genes), second for interaction with metabolites that contribute to TME-mediated immunosuppression (resulted in 17 genes), and third for the feasibility to detect relevant metabolites that are associated with the gene products (resulted in 11 genes) These 11 candidate genes were then experimentally filtered for efficient CRISPR/Cas9-based gene disruption in HEK293FT cells then CD8 T cells (resulted in 5 genes), for the feasibility to detect associated metabolites via LC/MS (resulted in 3 genes), and finally, for their functional importance, resulting in two candidates that passed all criteria: *ADA* and *PDK1*.

**Fig. 1:**
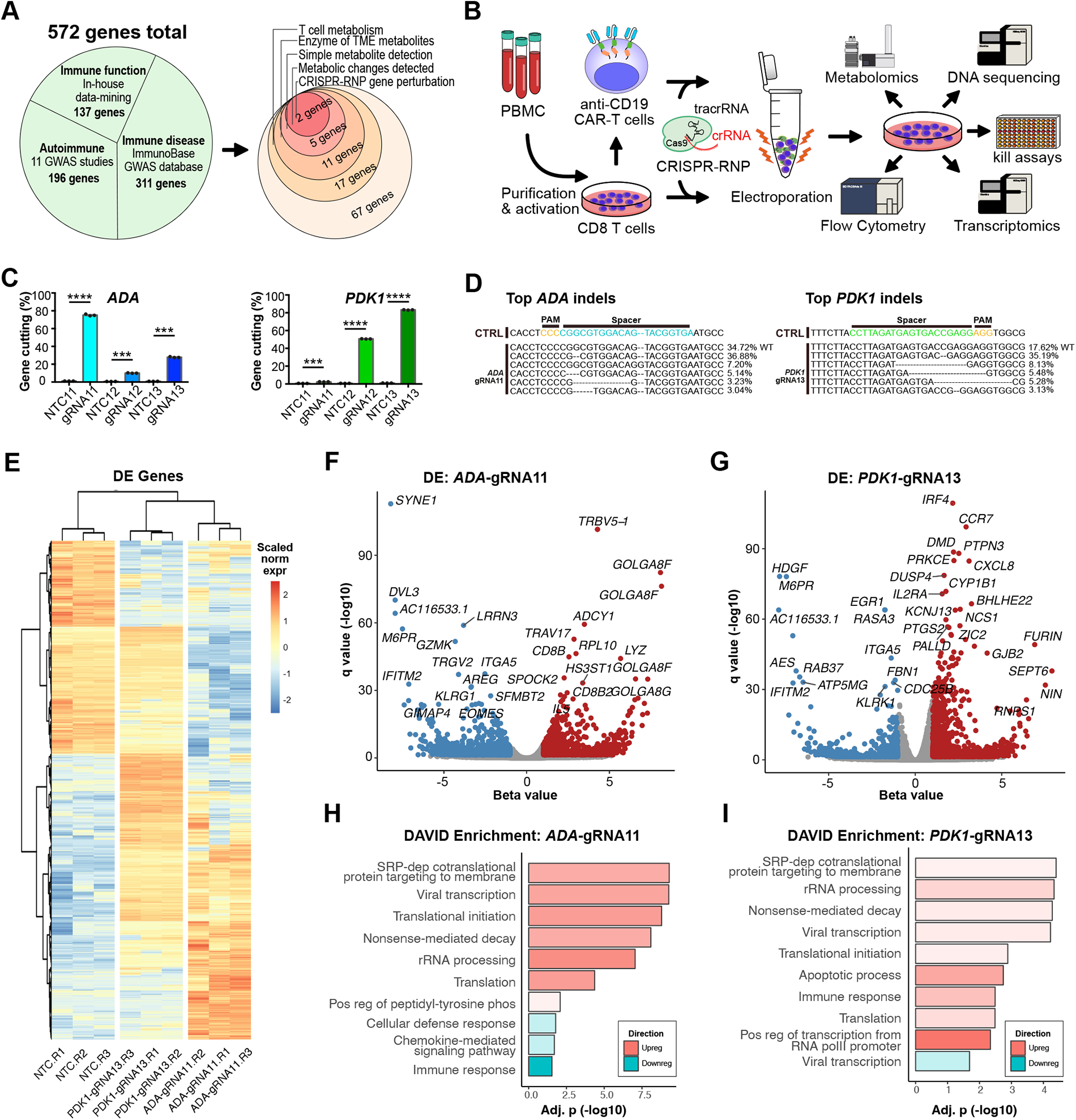
*In silico* screen and mRNA-seq revealed *ADA* and *PDK1* as metabolic reprogramming genes that induce immune response-related transcriptomic changes in CD8 T cells. **A,** Flowchart of the multi-tier selection for identification of candidate genes that modulate T cell metabolism. **B,** Schematic of mutant T cell and CAR T cell generation, multi-omics profiling and functional analyses. **C-D,** Quantification of CRISPR-mediated indel formation to *ADA* and *PDK1* genes. Sample sequences and indel statistics are displayed in C and D, respectively. **E,** Heatmap of significant differentially expressed (DE) transcripts in *ADA*-gRNA11 or *PDK1*-gRNA13 T cells. **F-G,** Volcano plots of differential expression (DE) in (F) *ADA*-gRNA11 or (G) *PDK1*-gRNA13 T cells. Expression fold-changes and significance are represented by the beta-value and q value, respectively. **H-I,** Waterfall plots of significantly enriched Biological Process ontologies in (H) *ADA*-gRNA11 or (I) *PDK1*-gRNA13 T cells. Results are shown for pathway analyses of DE genes, and significance is provided as the Benjamini-adjusted p value (adj. p). Enrichment was assessed separately for up- and down-regulated genes, but results are shown together. DE genes had a beta-change > 1 and q < 0.05. Indel quantification and DE were analyzed with 3 biological replicates for each condition.

### Generation of ADA and PDK1 knockout and overexpression mutant human CD8 T cells

To assess how *ADA* and *PDK1* genes impact the function of CD8 T cells, we generated target gene overexpression (OE) mutants using lentiviral vectors, and knockout (KO) mutants via electroporation of recombinant crRNA:tracrRNA/Cas9 ribonucleoprotein (CRISPR-RNP) into primary human CD8 T cells (**Fig. 1B**). Efficient gene editing was observed in the mutants, based on Nextera-based amplicon capture – next generation sequencing (NGS) of the target locus (**Fig. 1C-D**; **Supplementary Data 2**).

### ADA and PDK1 mutant CD8 T cells showed broad transcriptomic and metabolomic alterations

To reveal the effect of altered *ADA* and *PDK1* expression, mRNA-seq analyses were performed for each of the CD8 T cell mutants (**Fig. 1E-G**; **Supplementary Data 3**). Our data confirmed the knock-out of *ADA* or *PDK1* in mutant T cells, as each biological replicate showing significant downregulation of their respective gRNA target gene **(Supplementary Fig. 1A-B**). In the *ADA* mutants, gene enrichment analyses in *ADA*-gRNA cells showed that upregulated genes were associated with transcriptional and translational processes, while downregulated genes in *ADA* mutants are significantly associated with immune response and chemokine-mediated signaling (**Fig. 1H-I** **and Supplementary Fig. 1C**; **Supplementary Data 3**). The *ADA*-gRNA also significantly downregulated cytotoxicity genes: *GZMA*, *PRF1*, *IFNG* and *TNF*; exhaustion genes: *CD244*, *VSIR* and *LAG3*; and upregulated *IRF4*, a key transcription factor (TF) for *PDCD-1* expression and T cell exhaustion (**Supplementary Fig. 1D-F**) [31]. Overall, the *ADA*-gRNA downregulated effector genes for cytotoxicity, extra-lymphoid homing and adhesion, while upregulating the exhaustion genes in CD8 T cells.

In the *PDK1*-gRNA CD8 T cells, the most significant transcription changes include upregulation of the *IRF4* exhaustion TF and *CCR7* lymphoid homing receptor (**Fig. 1G**; **Supplementary Data 3**). There is a significant upregulation of *IFNG* and anti-inflammatory *IL10* cytokines, as well as upregulation of genes for common memory T cell markers, *CCR7* and *IL7R*, yet there is also a strong downregulation of core memory-related transcriptional regulator genes: *TCF7* and *BCL6* (**Supplementary Fig. 1E**; **Supplementary Data 3**). In addition, there is upregulation of multiple transcripts of *MYB*, a key memory transcription factor that activates *TCF7* transcription [32] (**Supplementary Fig. 2B**). There were transcriptional changes to immune checkpoint genes, including an upregulation of *LAG3* and *CD244* (**Supplementary Fig. 1F**). Gene ontology analyses shows enrichment in several transcriptional and translational ontologies, where the upregulated genes were enriched for biological processes, including immune response, inflammatory response, and the negative regulation of cell proliferation (**Fig. 1I** **and Supplementary Fig. 1C**; **Supplementary Data 3**).

To further understand the metabolic impact of the *ADA* and *PDK1* mutations in the CD8 T cells, we performed metabolomics profiling. The *ADA*-gRNA T cells showed the strongest metabolite shifts in purine metabolites, including a significant increase in GMP, ADP, adenosine (Ado), and deoxyadenosine (dAdo), as well as the purine biosynthesis intermediate PRPP (**Fig. 2A** **and Supplementary Fig. 2A**; **Supplementary Data 4**). Purine metabolism was also the most enriched pathway in a quantitative metabolic pathway analysis (MetPA, MetaboAnalyst v4.0) that integrates both metabolomics and transcriptomics data [30,33] (**Fig. 2C** **and Supplementary Fig. 2C**; **Supplementary Data 4**), and the pathway favors purine biosynthesis over salvage or catabolism with increased PRPP, XMP, GMP, S-AMP, and ADP, while no significant changes were found in hypoxanthine and IMP levels. Overexpression of *ADA* led to a significant increase in glucose, inosine and isoleucine, while there is a significant decrease in histidine and adenosine, the substrate of ADA (**Fig. 2A** **and Supplementary Fig. 2A**). MetPA analysis showed a significant association with the purine metabolism pathway with changes to levels of several metabolic intermediates and significantly increased inosine levels (**Fig. 2E** **and Supplementary Fig. 2D; Supplementary Data 4**). These data suggest that *ADA*-gRNA increased purine biosynthesis, and overexpression of *ADA* primarily affected the metabolites directly related to the ADA enzymatic reaction.

**Fig. 2:**
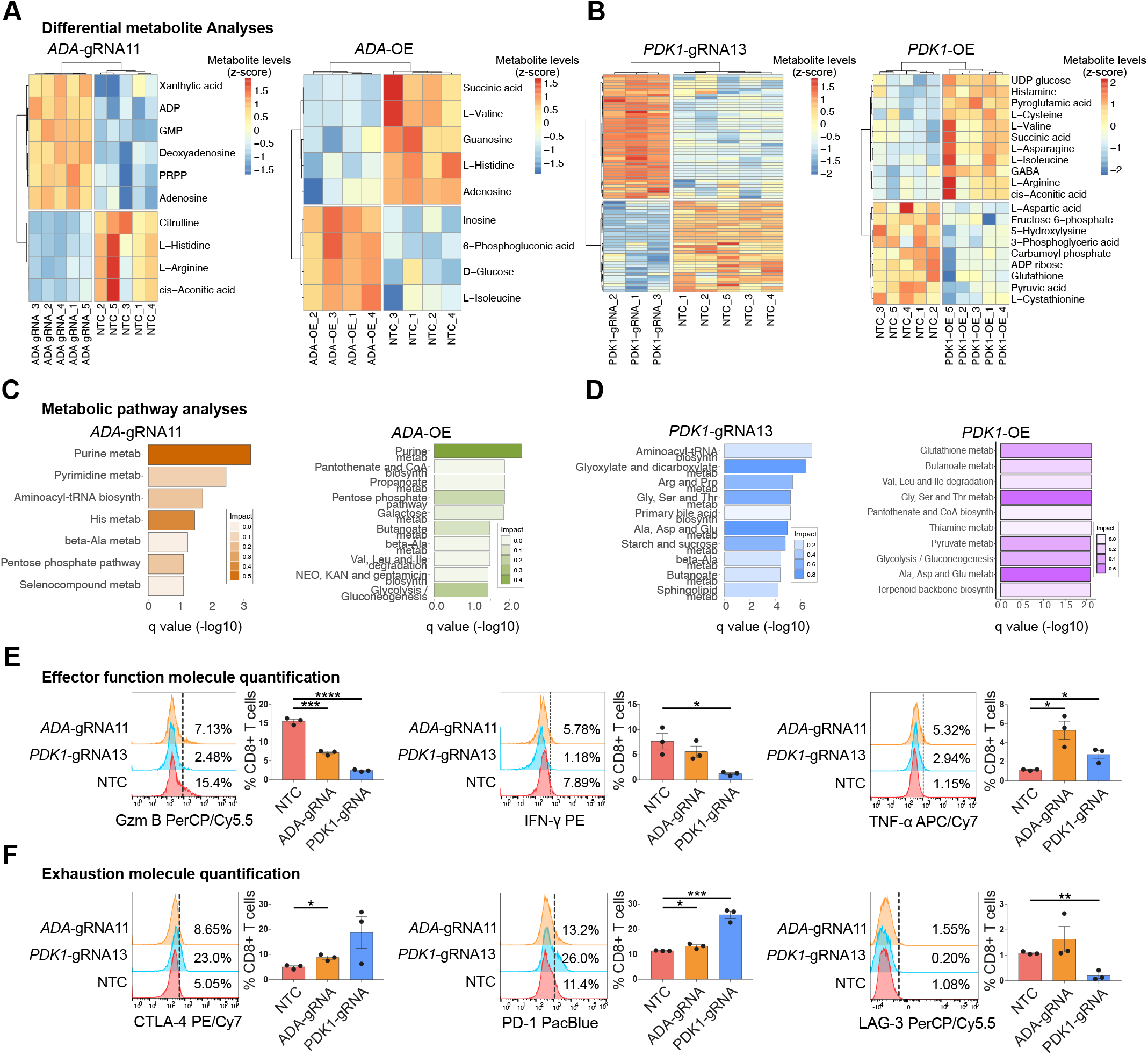
Metabolic reprogramming in *ADA* and *PDK1* mutant human primary T cells. **A-B,** Heatmaps of differential metabolite levels in (**A**) *ADA* and (**B**) *PDK1* T cell mutants. Significant metabolite changes had an adjusted-p < 0.10 and log2 fold-changes > 0.50. Heatmap values depict the z-score of log2 fold-changes. **C-D,** Waterfall plots of metabolic pathway analyses (MetPA) in (**C**) *ADA* and (**D**) *PDK1* T cell mutants. Bar color, size and opacity represent pathway significance and impact scores, which summarizing metabolite fold-changes of each pathway. MetPA results are given for metabolites-only in the overexpression mutants and for integrated metabolite-gene expression in the knockout mutants. **E-F,** Flow cytometric analyses of functional markers in *ADA* and *PDK1* T cell mutants. Histograms and are provided for (**E**) cytotoxicity and (**F**) exhaustion markers. Metabolomics and flow cytometry data were analyzed with n >= 3 for each condition and control.

Knocking out the *PDK1* significantly affected the levels of 71 metabolites in CD8 T cells, most prominently were several amino acids (**Fig. 2B** **and Supplementary Fig. 2B**; **Supplementary Data 4**). The integrated MetPA of mRNA-seq and metabolomics data showed significant association with aminoacyl-tRNA biosynthesis and the glutathione metabolism pathway, which showed significant upregulation of the glutathione pathway enzyme genes, as well as glutathione (**Fig. 2D** **and Supplementary Fig. 2E**; **Supplementary Data 4**), which has key roles in T cell function, including as a cellular anti-oxidant and modulating glycolysis and glutaminolysis through regulation of mTOR and NFAT [34]. Overexpression of *PDK1* significantly increased the levels of pyroglutamic acid and L-asparagine, while pathway analyses showed significant changes to glutathione and butanoate metabolism (**Fig. 2B, D and Supplementary Fig. 2B, E-F**; **Supplementary Data 4**). Together, we showed that in primary human CD8 T cells, *ADA* modulated metabolism through the purine biosynthesis pathway and influenced effector function genes, including *TNFA*, *IFNG*, granzymes and *PRF1*. In contrast, *PDK1* levels corresponded to changes in broad range of metabolic pathways, most notably glutaminolysis, glutathione biosynthesis, and the TCA cycle. In addition, the knockout of *PDK1* led to a significant increase of both effector and memory T cell genes. We assessed immune functions in the T cell mutants via flow cytometry to show a shift of both the *ADA*-gRNA and *PDK1*-gRNA T cells in the production of cytotoxic molecules with decreased granzyme B and IGN-γ and increased TNF-α (**Fig. 2E** **and Supplementary Fig. 3A**; **Supplementary Data 8**). In addition, there were increased levels of exhaustion markers in both mutant T cells, based on the significantly increased expression of CTLA-4 and PD-1 in *ADA*-gRNA, and of PD-1 and LAG-3 in *PDK1*-gRNA CD8 T cells (**Fig. 2F** **and Supplementary Fig. 3A-B**; **Supplementary Data 8**).

### ADA and PDK1 modulation improve cytotoxicity and survival in CD8 CAR-T cells

Modulating the expression of *ADA* and *PDK1* were shown to have strong effects to T cell metabolism, as well as cytotoxicity and survival gene expression. We next sought to test these functional attributes with a cytotoxicity assay by co-culture of T cells and cognate cancer cells. We generated CAR-T cells using a lentiviral vector with an anti-CD19 CAR (FLAG-(anti-CD19 scFv)-CD137-CD3ζ) [9,10]. The *ADA* or *PDK1* gene was constitutively overexpressed (OE) on the same lentiviral CAR-T vector under a T2A (self-cleaving peptide) element (**Fig. 3A**). The overexpression or control CAR-T cells were then cultured and sequentially FACS-sorted for CAR-T purity (**Fig. 3B** **and Supplementary Fig. 3C**), and then *ADA/PDK1*-gRNA cells were generated from the control CAR-T cells via CRISPR-RNP mediated knockout (**Fig. 1B, 3A-B**), with efficient gene perturbations detected by Nextera-based amplicon sequencing and western blots (**Fig. 3C-E**; **Supplementary Data 2**). The different T cell populations were then co-cultured with NALM6-GL B cell precursor leukemia cell line constitutively expressing GFP and firefly luciferase [9] at different concentrations to evaluate T cell-mediated cancer killing capacity. Our results showed that the overexpression of either *ADA* or *PDK1* significantly enhanced cancer cytolysis by the CAR-T cells, based on NALM6-GL luciferase bioluminescence (**Fig. 3F**; **Supplementary Data 5**). The *ADA*-OE cytolysis consistently increased at all cancer cell concentrations (titrated as effector:target (E:T) ratios), whereas the *PDK1*-OE cytolysis is most significantly increased at intermediate cancer cell concentrations and non-significantly changed at high concentrations. In contrast, both the *ADA*-gRNA and *PDK1*-gRNA CAR-T cells have decreased cytolytic capabilities at all cancer cell concentrations.

**Fig. 3:**
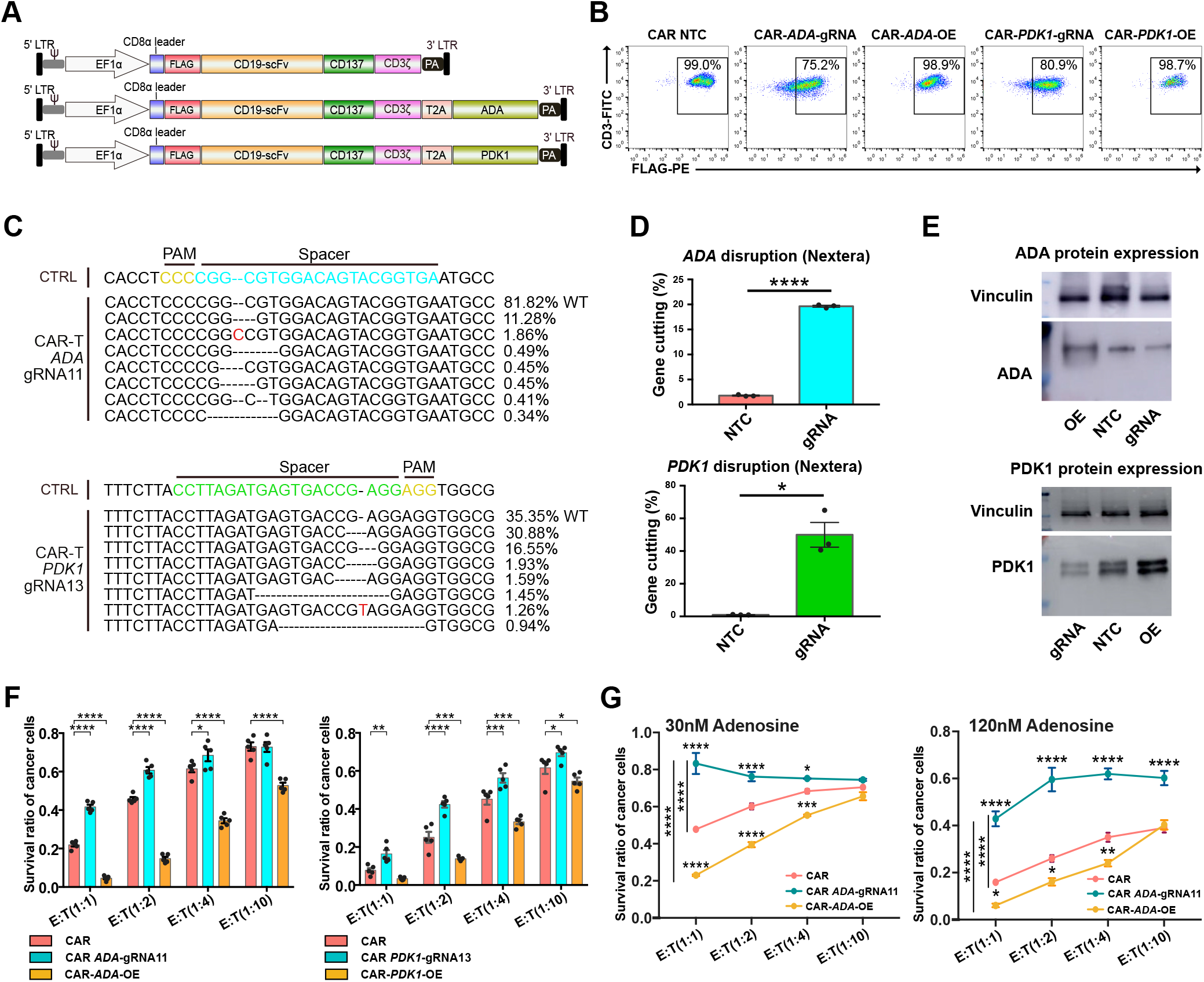
Overexpression of *ADA* or *PDK1* in α-CD19 CAR-T cells enhanced cytotoxicity. **A,** Schematic maps of the α-CD19 CAR, CAR-*ADA*-OE, and CAR-*PDK1*-OE lentiviral vectors. **B,** Flow cytometry dot plots of α-CD19 CAR protein expression in CAR-T cell populations from a representative sample. **C-E,** CRISPR-mediated perturbations to *ADA* and *PDK1* gene loci in CAR-T cell populations. Indel formation and protein expression were assessed by (**C-D**) Nextera DNA sequencing and (**E**) Immunoblot, respectively. **F-G,** Bar plots and line plots of co-culture cytotoxicity assay results for different CAR-T cell populations. Cytotoxicity was quantified as NALM6-GL cancer cell survival in co-culture assays with CAR-T cells at different effector-target ratio concentrations (E:T). NALM6-GL levels were quantified by luciferase bioluminescence, relative to no treatment. Results are shown for (**F**) *ADA*/*PDK1* vs control CAR-T cells via Dunnett’s two-way ANOVA test, and the (**G**) *ADA* vs control CAR-T cells in 30 nM and 120 nM adenosine. All analyses were performed with n = 3 biological replicates for each condition.

We next assessed whether *ADA*-OE increased cancer killing by decreasing the extracellular Ado levels, which are immunosuppressive when increased in the tumor microenvironment [35]. To address this, cancer cytolysis was assessed in elevated and high Ado concentrations (30uM and 120uM, respectively) [36–38]. *ADA*-OE CAR-T cells showed enhanced killing overall and at most E:T ratios, while *ADA*-gRNA decreases killing in the presence of Ado (**Fig. 3G**; **Supplementary Data 5**). Overall, these data showed that overexpressing either *ADA* or *PDK1* leads to enhanced cancer cell cytolysis for anti-CD19 CAR-T cells, while the *ADA*-OE also enables increased cytotoxicity under high concentrations of extracellular Ado.

### Transcriptomic and metabolomic alterations in ADA-OE and PDK1-OE CAR-T cells

To determine how modulation of *ADA* and *PDK1* affect T cells with the CAR system, we performed mRNA-seq transcriptional profiling of the different CAR mutants in parallel with matched anti-CD19 CD8 CAR-T controls (**Fig. 4A-B, D**; **Supplementary Data 6**). Differential expression (DE) of *ADA*-gRNA CAR-T cells showed upregulated *GZMB* and *GZMM,* as well as the *BCL2* pro-survival gene, while pathway analyses upregulated genes in the type-I interferon, TNF-alpha and IFN-gamma response pathways (**Fig. 4C** **and Supplementary Fig. 4A & 5A, D-F**). These combined data showed an upregulation of cytotoxicity, survival and lymphoid trafficking factors. The expression profile in *ADA*-OE CAR-T cells also showed a cytotoxic profile with a significant upregulation of *GZMA, GZMB* and *GZMM* and downregulation of memory-related transcription factor genes: *STAT3*, *HDAC5* and *FOXO1* genes, as well as a significant switch in *PRDM1* isoforms [39] (**Supplementary Fig. 4A**; **Supplementary Data 6**). Pathway analyses showed significantly upregulated genes are enriched in type-I IFN signaling and chemotaxis pathway (DAVID analyses), as well as less significant IFN-α and IFN-γ response upregulation (GSEA analyses) (**Fig. 4E** **and Supplementary Fig. 5B, D-F**; **Supplementary Data 6**). Together, these data showed an increased cytotoxic signature with altered memory transcriptional regulatory profile.

**Fig. 4:**
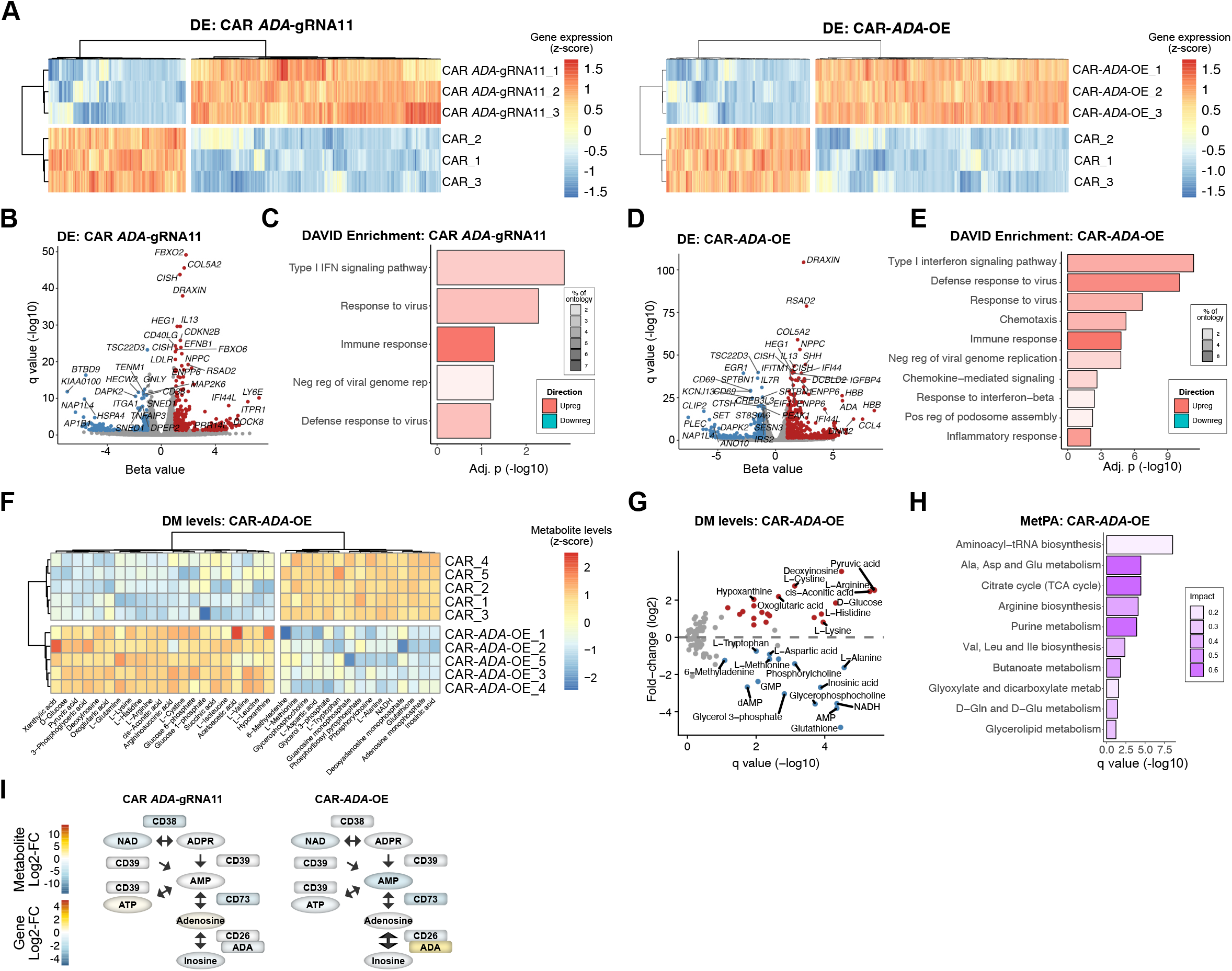
Multi-omics profiling and phenotypic characterization of *ADA* mutant CAR-T cells. **A,** Heatmaps of significant differentially expressed (DE) transcripts in *ADA*-gRNA11 and *ADA*-OE CAR-T cell mutants. **B, D,** Volcano plots of DE analyses in (**B**) *ADA*-gRNA11 and (**D**) *ADA*-OE CAR-T cells. Expression fold-changes and significance are represented by the beta-value and q value, respectively. **C, E,** Waterfall plots of significantly enriched Biological Process ontologies in (**C**) *ADA*-gRNA11 and (**E**) *ADA*-OE CAR-T cells. Results are shown for DAVID analyses of DE genes, and significance is provided as Benjamini-adjusted p values (adj. p). Enrichment was assessed separately for up- and downregulated genes, but results are shown together. **F,** Heatmap of Significant differential metabolite (DM) levels in *ADA*-OE CAR-T cell mutants. **G,** Volcano plot of differential metabolite levels in *ADA*-OE CAR-T cell mutants. **H,** Waterfall plot of integrated transcriptomic-metabolomic MetPA in *ADA*-OE CAR-T cells. Bar color, size and opacity represent pathway significance and impact scores, which summarizing metabolite fold-changes of each pathway. **I,** Partial pathway map of extracellular adenosine metabolism in *ADA* CAR-T cell mutants. Log2-fold-changes of normalized metabolites and enzyme genes are depicted by ovals and rectangles, respectively (heatmap color-scale). Multi-gene enzyme levels are presented as log2-fold-changes of the sum of normalized gene counts. Metabolomics and transcriptomics data were analyzed with n = 5 and n = 3 biological replicates, respectively, for each condition. DE genes had a beta-change > 1 and q < 0.05, while DM had adjusted-p < 0.10 and a log2 fold-change > 0.50. Volcano plot labels are given to genes/metabolites with the highest significance and fold-changes, for which significant increases and decreases are shown in red and blue, respectively.

Metabolic profiling of the *ADA*-OE CD8 CAR-T cells showed significantly altered the levels of 37 metabolites, including a strong increase in pyruvate and several TCA intermediates (cis-Aconitate, a-ketoglutarate, succinate), as well as a significant increase in glutamine and a decrease in glutamate (**Fig. 4F-G**; **Supplementary Data 7**). The RNA-Metabolite integrated MetPA indicated significant changes to several aminoacyl and aminoacyl-tRNA biosynthesis pathways (**Fig. 4H** **and Supplementary Fig. 7A**; **Supplementary Data 7**). The enrichment analyses also found associations with purine metabolism, upregulation of the TCA cycle with significantly upregulated intermediates, and downregulation of glutathione metabolism (**Fig. 4H-I** **and Supplementary Fig. 7A**). Overall, the metabolomics of *ADA*-OE CAR-T cells showed changes to several aminoacyl metabolic pathways, suggesting an increased usage of glutaminolysis and at least partial TCA usage, in which the synthesis of intermediates α-KG and succinate were disproportionately promoted. The purine pathway was marked by an upregulation of multiple intermediates (cAMP, adenylsuccinate, dATP), as well as both increased adenosine and decreased inosine (**Supplementary Data 7**), which was expected by a decrease of the ADA enzyme. There was also increased Ado, AMP and ATP in *ADA*-gRNA relative to *ADA*-OE CAR-T cells (**Fig. 4I; Supplementary Data 7**). Together, altering the expression of *ADA* still predominantly affected purine metabolism, where the *ADA*-OE CD8 CAR-T cells showed metabolic changes, which were marked by increased glutaminolysis and altered TCA usage.

In the differential expression (DE) analysis of *PDK1*-OE CD8 CAR-T cells, there was a significantly increased expression of multiple cytotoxicity genes, including *GZMA*, *GZMB*, *PRF1*, *LAMP3*, and *IFNG*, all indicating increased effector functions (**Fig. 5A-B** **and Supplementary Fig. 4B**; **Supplementary Data 6**). There were also increased expression of exhaustion-related checkpoint genes, such as *TIGIT*, *CD244* and *LAG3*, as well as upregulation of *EOMES* and *IRF4* transcription factors, and increased expression of antiapoptotic gene *BCL2* (**Supplementary Fig. 4B**). Pathway analyses showed that upregulated genes were significantly enriched in type-I IFN and IFN-γ responses, cell-cell signaling and chemotaxis (DAVID analyses), as well as the IFN-alpha and IFN-gamma response, with decrease in the expression of MYC targets (**Fig. 5C** **and Supplementary Fig. 5C & 6A-C**; **Supplementary Data 6**). Together, these data suggested a strong effector phenotype with a notable increase in pro-survival genes, yet T cell exhaustion markers were elevated. In contrast to this expression profile, the *PDK1*-gRNA CAR-T cells displayed no significant changes to effector function genes (**Supplementary Fig. 4B**; **Supplementary Data 6**). There was a significant increased expression of pro-survival *BCL2* and strong decreased expression of the *MYB* memory regulatory gene (**Supplementary Fig. 4B**).

**Fig. 5:**
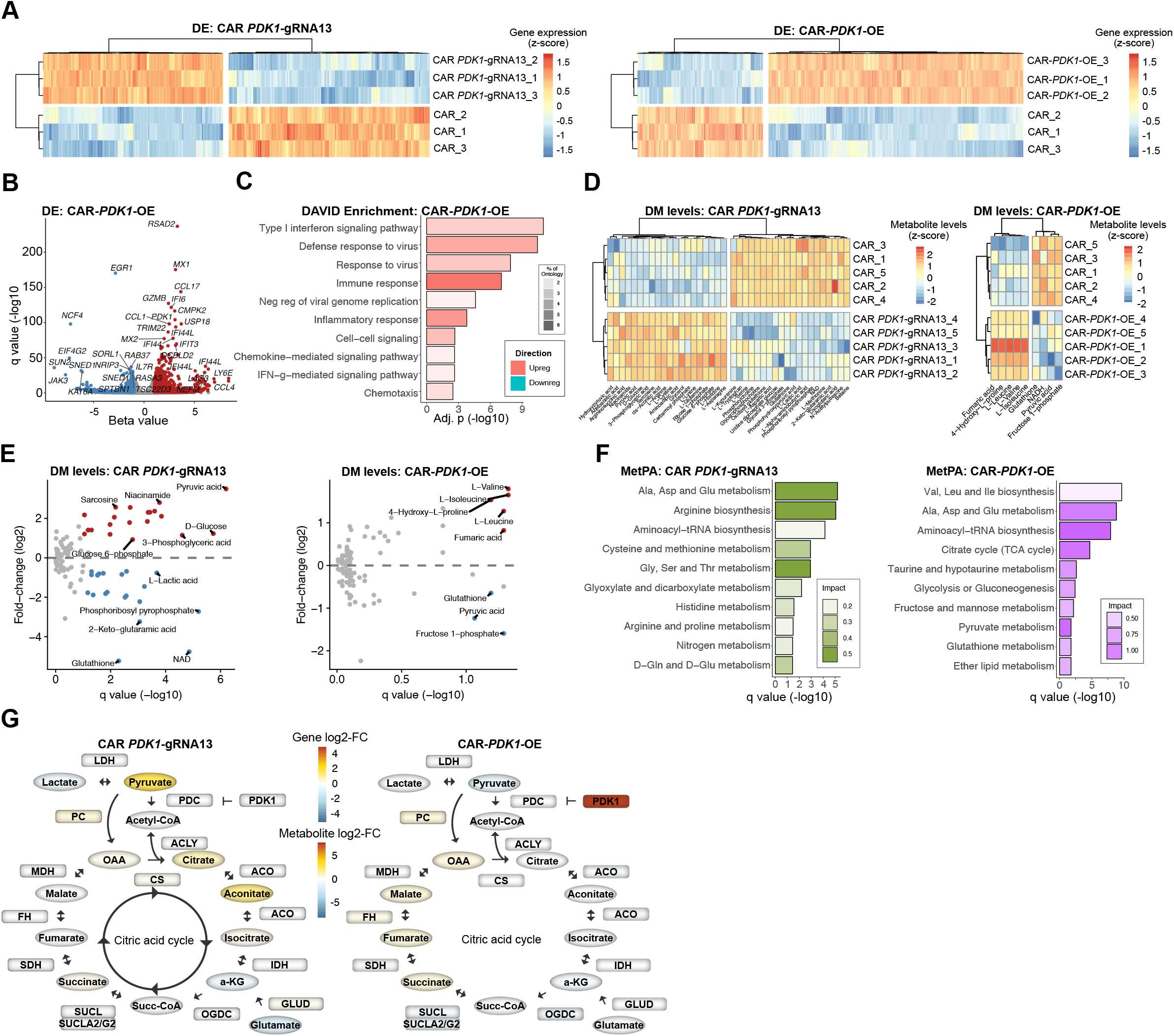
Multi-omics profiling and phenotypic characterization of *PDK1* mutant CAR-T cells. **A,** Heatmaps of significant differentially expressed (DE) transcripts in *PDK1*-gRNA13 and *PDK1*-OE CAR-T cell mutants. **B,** Volcano plot of DE analyses in *PDK1*-OE CAR-T cells. Expression fold-changes and significance are represented by the beta-value and q value, respectively. **C,** Waterfall plots of significantly enriched Biological Process ontologies in *PDK1*-OE CAR-T cells. Results are shown for DAVID analyses of DE genes, and significance is provided as the Benjamini-adjusted p value (adj. p). Enrichment was assessed separately for up- and downregulated genes, but results are shown together. **D,** Heatmaps of differential metabolite (DM) levels in *PDK1*-gRNA13 and *PDK1*-OE CAR-T cells. **E,** Volcano plot of DM levels in *PDK1*-gRNA13 and *PDK1*-OE CAR-T cells. **F,** Waterfall plots of integrated transcriptomic-metabolomic MetPA in *PDK1*-gRNA13 and *PDK1*-OE CAR-T cells. Bar color, size and opacity represent pathway significance and impact scores, which summarizing metabolite fold-changes of each pathway. **G,** Partial pathway maps of the TCA cycle in *PDK1*-gRNA13 and *PDK1*-OE CAR-T cell mutants. Log2 fold-changes of metabolite and gene levels are depicted by ovals and rectangles, respectively (heatmap color-scale). Multi-gene enzyme levels were calculated as the log2-fold-changes of the sum of normalized gene counts. Metabolomics and transcriptomics data were analyzed with n = 5 and n = 3 samples, respectively, for each condition and control. DE genes had a beta-change > 1 and q < 0.05, while DM had adjusted-p < 0.10 and a log2 fold-change > 0.50. Volcano plot labels are given to genes/metabolites with the highest significance and fold-changes, for which significant increases and decreases are shown in red and blue, respectively.

For the metabolomics of *PDK1*-OE CD8 CAR-T cells, the most prominent significant changes were increases in valine and isoleucine (**Fig. 5D-E**; **Supplementary Data 7**). Integrated pathway analysis showed significant associations with several metabolic pathways, including multiple amino acid metabolic pathways and the TCA cycle (**Fig. 5F** **and Supplementary Fig. 7C**; **Supplementary Data 7**). The amino acid changes were most pronounced in the neutral alkyl residues: valine, isoleucine and leucine, which were all increased, while the TCA cycle changes reflect a subtle increase in TCA utilization (**Fig. 5F-G** **and Supplementary Fig. 7C**). The *PDK1*-gRNA CAR-T cells had significant changes up-to 36 metabolites (q < 0.05), with the most extreme changes including an upregulation of pyruvate and glucose and a downregulation of PRPP and nicotinamide adenine dinucleotide (NAD) (**Fig. 5D-E**; **Supplementary Data 7**). There was also a noteworthy significant decrease in the level of lactate (**Supplementary Data 7**). The MetPA showed significant associations with several amino acid metabolic pathways, particularly the alanine-aspartate-glutamate metabolism (decreased amino acid pathway products) and arginine biosynthesis pathway (**Fig. 5F** **and Supplementary Fig. 7B**; **Supplementary Data 7**). Overall, the *PDK1*-gRNA indicated upregulation of glycolysis and early TCA intermediates with a strong depletion of NAD.

### PDK1-OE reduces stability of CAR surface expression and cytotoxicity over time

Given the relationship between metabolism and T cell function, we sought to investigate how *ADA*-OE and *PDK1*-OE influenced the T cell phenotype. Therefore, we performed multi-faceted functional characterizations of *ADA*-OE and *PDK1*-OE CD8 CAR-T cells by flow cytometry. First, we evaluated effector T cell function in stimulated CAR-T cells (4 hours), which showed unexpected decreases in the early production of granzyme B, perforin, IFN-γ, and TNF-α with both *ADA*-OE and *PDK1*-OE (**Supplementary Fig. 8A**). As decreased effector function is counterintuitive to the increased cancer cell lysis from our competition assays (**Fig. 3F**), we performed another competition assay that showed ADA-OE still had improved cancer cell lysis, but *PDK1*-OE CAR-T cells had lost its cytotoxicity (**Supplementary Fig. 8B**). Further analysis demonstrated that the surface expression of the CAR was lost in PDK1-OE, but not ADA-OE or control cells (**Supplementary Fig. 8C**). These experiments were performed using copy-number-controlled knock-in/knock-out (KIKO) CAR system, in which CRISPR-RNP was electroporated into CD8 T cells to induce dsDNA breaks in the *TRAC* locus, followed by cell infection with an adeno-associated virus type 6 (AAV6) vector with anti-CD19 CAR, CAR-T2A-*ADA*, or CAR-T2A-*PDK1* [9]. The progressive downregulation of CAR in PDK1-OE T cells would explain the discrepancy between the competition assay results in **Fig. 3F** and **Supplementary Fig. 8C**, which were respectively performed days and weeks after FACS selection of CD8+CAR+ T cells. The remaining experiments were performed in KIKO CAR-T cells to exclude the potential influence of *ADA*/*PDK1* copy-number inconsistencies from using lentiviral vectors. In addition, CAR-T cells were sorted only once for CD8+CAR+CD3-cells to prevent result biases from gradual CAR downregulation.

### Overexpression of ADA increases cytotoxicity and memory, while decreasing exhaustion in CAR-T cells

To further explore how *ADA*-OE CAR-T cells have enhanced cancer killing even though the production of effector molecules is initially low (**Supplementary Fig. 8A**), we assessed effector function and cancer lysis over multiple time points. Our results revealed that the *ADA*-OE cells significantly increased granzyme B production at 2- and 4-days post stimulation (**Fig. 6A**), coinciding with enhanced cancer cell clearance (**Fig. 6B**). Immune profiling also demonstrated that *ADA*-OE also enhances proliferation upon stimulation, based on assays with CellTrace Violet, which involves labeling cells with a fluorescent dye whose concentration decreases across subsequent cellular divisions. These assays demonstrated that nearly all *ADA*-OE CAR-T cells undergo at least one round of division (**Fig. 6C**). In addition, *ADA*-OE increased polarization of central memory (TCM; CD45RO+CD62L+) vs effector memory T cell subsets (TEM; CD45RO+CD62L-) (**Fig. 6D**). Next, we profiled markers of T cell exhaustion, showing that *ADA*-OE decreases LAG-3 and TIM-3, but not PD-1 and CTLA-4 (**Fig. 6E, Supplementary Data 8**). These data further revealed the immunological phenotype of *ADA* overexpression in CD8 CAR-T cells, which enhances cytotoxicity and memory, while decreasing exhaustion.

**Fig. 6:**
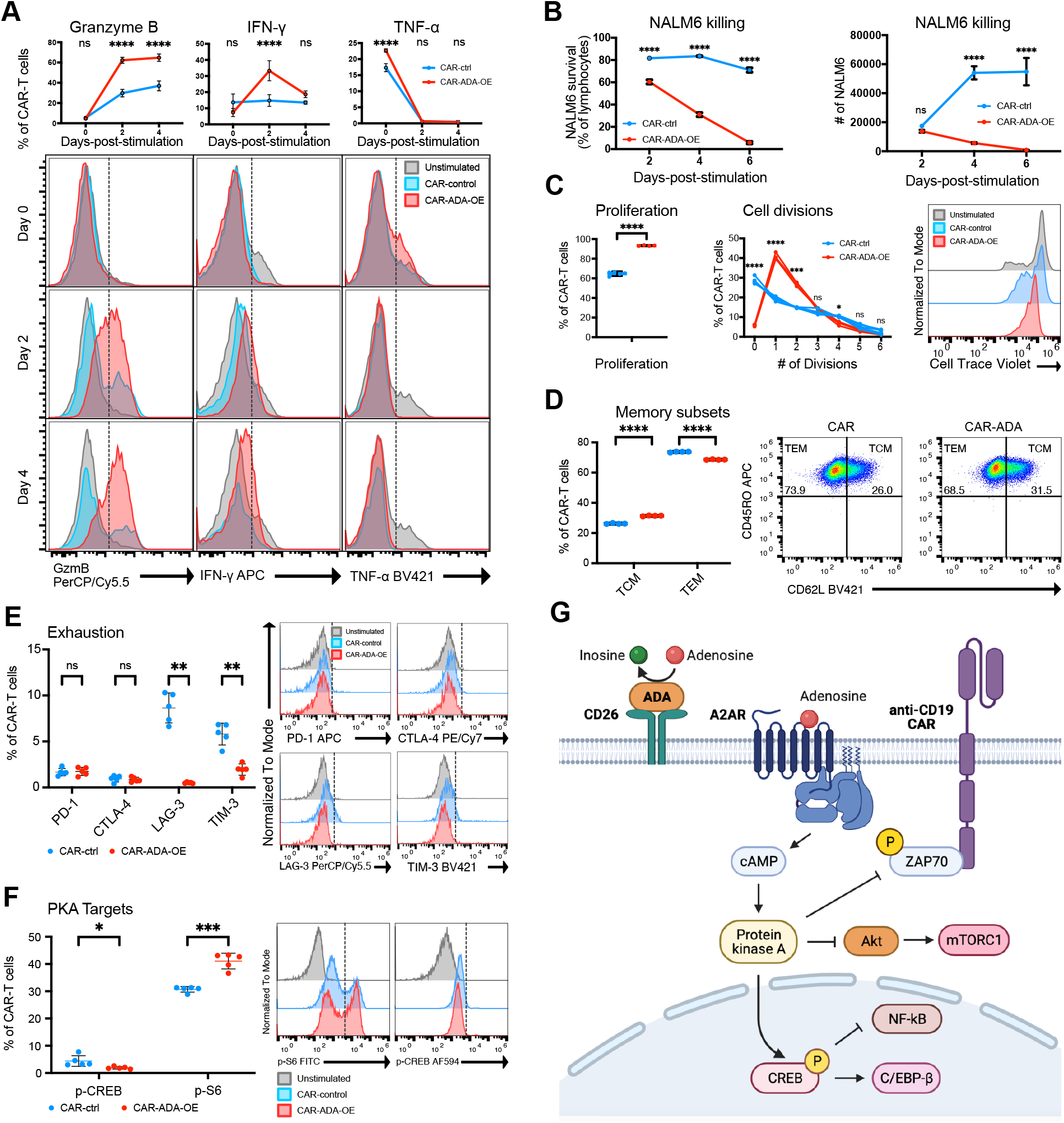
Phenotypic profiling of *ADA*-OE CAR-T cells. **A-B,** Flow cytometric analyses for effector molecule production and NALM6-GL cytolysis in *ADA*-OE vs control CAR-T cells at 0, 2, and 4 days post-stimulation in a co-culture assay. Cancer lysis was measured as GFP+ cell numbers or percent of live lymphocytes. **C,** Proliferation assay of stimulated CAR-T cells, measured as the dissipation of Cell Trace Violet dye across cell divisions (modeled by FlowJo software). Proliferation is compared at 0 vs any divisions and across each number of divisions. **D-F,** Flow cytometry analyses of markers for memory, exhaustion, and downstream targets of A2AR-PKA signaling. **G,** Schematic of A2AR-PKA signaling pathway. Experiments in (**A-F**) were performed with >= 4 biological replicates in >= 2 independent experiments. When indicated, CAR-T cells were stimulated with NALM6-GL at a 1:2 E:T ratio.

The mechanism of ADA should reduce extracellular adenosine, which we expect would decrease signaling by the adenosine receptor, and thus, should decrease PKA activation [40]. As PKA signaling in this context could lead to exhaustion through CREB activation and mTORC1 inhibition (**Fig. 6G**) [40], we assessed the downstream targets of PKA to show that ADA-OE CAR-T cells have significantly increased mTORC1 signaling and significantly decreased CREB activation, based on p-S6 and p-CREB levels, respectively (**Fig. 6F, Supplementary Data 8**).

### ADA Overexpression increases α-HER2 CAR-T cell function in colorectal cancer in vivo models

We set out to test ADA-OE in CAR-T cells generated from heterogeneous CD3+ T cell populations, and test them in an α-HER2 CAR *in vivo* solid tumor models. First, we knocked-in α-HER2 CAR construct to the *TRAC* locus using Cas9-RNP (**Fig. 7A**). The *in vitro* characterization assays were performed as above to demonstrate that ADA-OE improves α-HER2 CAR T similarly to α-CD19 CAR, including increased cytotoxicity (IFN-γ, granzyme B, TNF-α), decreased pCREB levels and exhaustion, marked by LAG-3 and PD-1 in both CD4 and CD8 CAR-T (**Supplementary Fig. S9A-D, Supplementary Data 8**). Next, ADA-OE was assessed in α-ΗΕR2 CAR-T cells that were adoptively transferred by intravenous injection to mice with subcutaneous HT29-GFP-Luciferase (HT29-GL) tumors (2 independent experiments of 4-5 mice per condition group). The ADA-OE CAR-T group showed enhanced anti-tumor efficacy compared to the control CAR-T and WT T cells (**Fig. 7B, Supplementary Data 8**). Tumors were extracted and analyzed, which showed that ADA-OE also increased CAR-T infiltration and CD8/CD4 CAR-T proportions, as well as proliferation rates and central memory polarization in both CD4 and CD8 populations (**Fig. 7C-E and Supplementary Fig. 10A-B, Supplementary Data 8**). ADA-OE also decreased the levels of PD-1 in CD8 CAR-T, but increased CTLA-4 in CD4 and CD8 CAR-T (**Supplementary Fig. 10C, Supplementary Data 8**). The remaining exhaustion markers were not significantly different between the treatment groups.

These results demonstrated that ADA-OE (1) increased the performance of α-HER2 CAR-T cells similarly to α-CD19 CAR-T *in vitro*; (2) enhanced function of CAR-T generated from CD3 T cells; (3) significantly improved anti-tumor efficacy of CAR-T treatment in an *in vivo* colorectal cancer model.

**Fig. 7:**
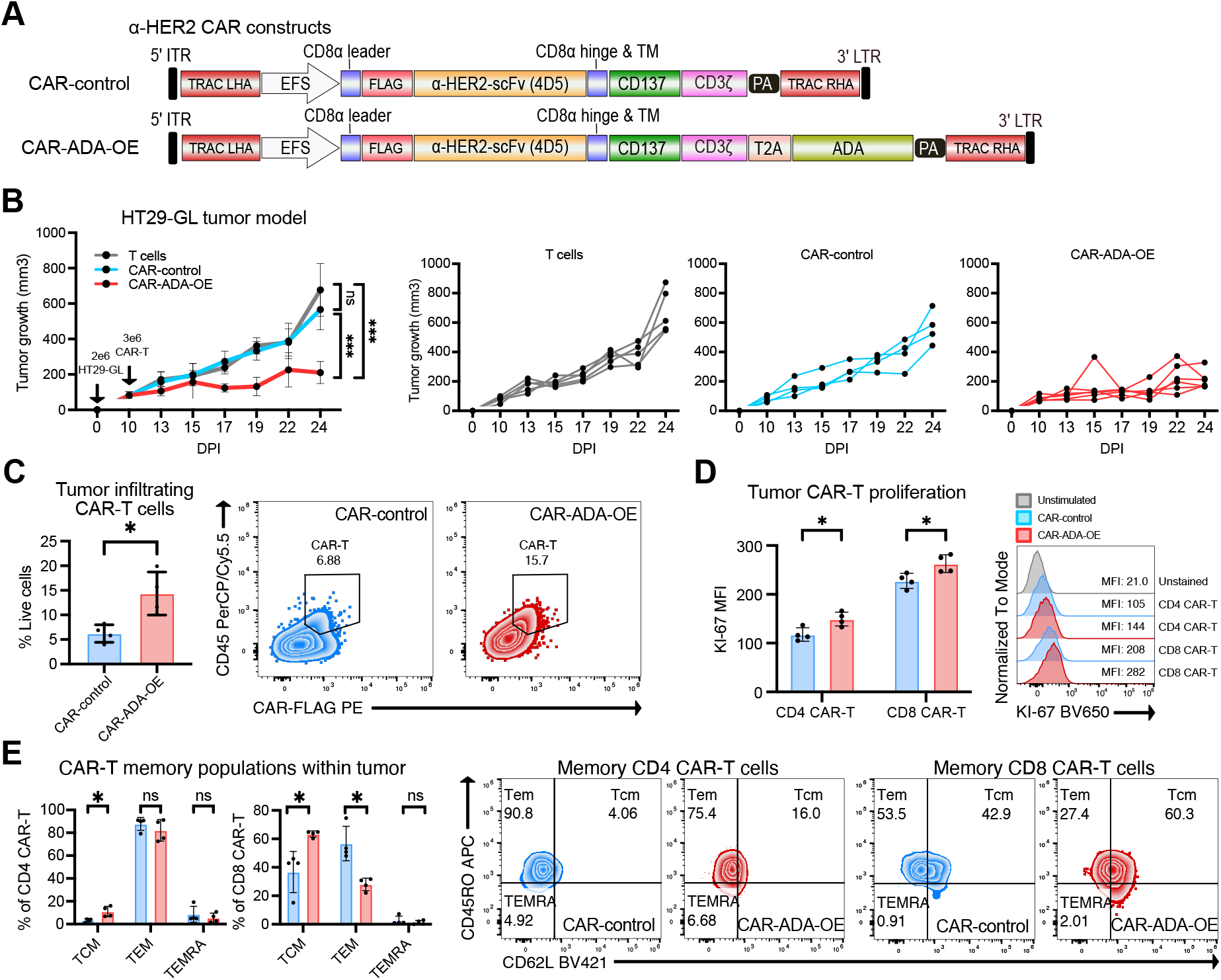
*ADA*-OE enhances α-HER2 CAR-T cell function in an *in vivo* colorectal cancer model. **A,** Schematic maps of the *ADA*-OE and control α-HER2 CAR lentiviral vector constructs. **B,** Tumor growth curves of *ADA*-OE and control α-HER2 CAR-T treatments in an *in vivo* HT29-GL tumor model. The left panel is shown with summary curves for each treatment, while the right panels show spider plot curves of the individual growth curves, separately graphed by treatment group. **C**, Flow cytometry analyses of the percentages of tumor infiltrating ADA-OE vs control α-HER2 CAR-T cells. Tumor infiltration percentage is relative to live, single cells. **D-E,** Flow cytometry analyses of ADA-OE vs control CAR-T (**D**) proliferation (KI-67) and (**E**) memory phenotypes in CD4 or CD8 CAR-T cells. Tumor growth and flow cytometry experiments included >= 4 mice per treatment with 2 independent experiments.

## Discussion

ADA catalyzes the conversion of adenosine to inosine intracellularly, as part of the purine metabolism pathways, and extracellularly, where ADA can co-stimulate T-cell activation through its interaction with CD26 and enhance both IFN-gamma and TNF-alpha production [41,42]. While intracellular adenosine is involved in energy metabolism, nucleic acid metabolism, and the methionine cycle [43], deficiency of *ADA* can lead to immunosuppressive effects through the accumulation of Ado and dAdo substrates. dAdo and dATP are released upon cell death and can induce apoptosis in T cells [44,45], whereas Ado and dAdo accumulation enhances adenosine receptor 2A (A2aR) signaling that disrupts T cell activation and function, including inhibition of mTORC and ERK1/2 signaling pathways, PKA hyperactivation, and increased intracellular cAMP, which can decrease IFN-gamma and IL-2 productions in T cells [40,46–49]. Similarly, we showed decreased cytotoxicity in *ADA*-deficient CAR-T cells and found significantly enhanced cytotoxicity and memory function in *ADA*-OE CAR-T cells. The cytotoxicity of *ADA*-OE is also enhanced under high concentrations of extracellular adenosine, an inhibitor of immune function that is greatly increased in the hypoxic conditions of the TME [50], where the adenosine can stimulating tumor growth and angiogenesis through fostering the tumor cells to material and energy metabolism [51,52].

PDK1 assists in the metabolic switch during T cell activation from oxidative phosphorylation to aerobic glycolysis, in which pyruvate is primarily converted to lactate for energy production [53]. This metabolic switch is bioenergetically favorable for rapid proliferation and cytokine production of effector T cells, and the early glycolytic shift is facilitated, in part, by PDK1 though inhibiting pyruvate decarboxylase and blocking pyruvate import to the mitochondria [54]. Our data showed that *PDK1* overexpression enhanced effector T cell function with the upregulation of cytokine, cytotoxicity and chemotaxis-related genes, and thereby improving CAR-T cell cytotoxicity. However, the *PDK1*-OE CAR-T cells are likely transitioning toward a terminally exhausted state, given the overexpression of genes related to T cell exhaustion and downstream targets of the TCF-1-exhaustion axis, in which TCF-1 drives expression of EOMES and c-Myb, that upregulates the anti-apoptotic Bcl-2 gene, ultimately leading to the development of a persistent exhausted T cell [55].

Taken together, we identified and omics-profiled ADA and PDK1 as key metabolic enzymes directly in human primary T cells and CAR-T cells. We demonstrated that ADA overexpression improves α-CD19 and α-HER2 CAR-T cells from different donors by enhancing cancer lysis, CAR-T proliferation and central memory generation, while decreasing exhaustion. Furthermore, our results revealed that ADA-OE significantly improves the efficacy of α-ΗΕR2 CAR-T cells in an *in vivo* solid tumor model. These findings highlight the roles of these new enzymes in the metabolism and T cell function critical for cancer immunotherapy and cell therapy.

## Acknowledgments

We thank Drs. Sznol, Fuchs, Bersenev, Isufi, Seropian, and Krause for discussion. We thank all members in Chen laboratory, as well as various colleagues in Department of Genetics, Systems Biology Institute, Cancer Systems Biology Center, MCGD Program, Immunobiology Program, BBS Program, Cancer Center, Stem Cell Center, Liver Center, RNA Center and Center for Biomedical Data Sciences at Yale for assistance and/or discussion. We thank the Center for Genome Analysis, Center for Molecular Discovery, High Performance Computing Center, West Campus Analytical Chemistry Core, West Campus Mass Spec Core and Keck Biotechnology Resource Laboratory at Yale, for technical support.

S.C. is supported by NIH/NCI/NIDA (DP2CA238295, R01CA231112, U54CA209992-8697, R33CA225498, RF1DA048811), DoD (W81XWH-17-1-0235, W81XWH-20-1-0072, W81XWH-21-1-0514), Damon Runyon Dale Frey Award (DFS-13-15), Melanoma Research Alliance (412806, 16-003524), St-Baldrick’s Foundation (426685), Breast Cancer Alliance, Cancer Research Institute (CLIP), AACR (17-20-01-CHEN), The Mary Kay Foundation (017-81), The V Foundation (V2017-022), Alliance for Cancer Gene Therapy, Sontag Foundation (DSA), Pershing Square Sohn Cancer Research Alliance, Dexter Lu, Ludwig Family Foundation, Blavatnik Family Foundation, and Chenevert Family Foundation. PR is supported by Yale PhD training grant from NIH (T32GM007499) and Lo Fellowship of Excellence of Stem Cell Research. JJP is supported by NIH MSTP training grant (T32GM007205). GW is supported by CRI Irvington and RJ Anderson Postdoctoral Fellowships. XD is supported by Charles H. Revson Senior Postdoctoral Fellowship.

## Contributions

SC conceived the study, and designed with PAR, QY, LY and JJP. JJP and QY performed the *in silico* screen. PAR, QY, and LY performed most experiments. In particular, PAR performed most immune studies, CAR-T construction and testing, all NGS data analysis, and *in vivo* tumor immunology analyses. QY and LY performed primary T cell analysis, CAR-T construction, gene KI/OE and KO, RNA-seq and cellular analyses. QY performed most metabolomics experiments. PAR and MB performed *in vivo* HER2 CAR-T solid tumor efficacy study. WHL, AA, DL, YZ, XD, GW, YE, TW and PC assisted certain experiments, or provided resources or conceptual inputs. SC secured funding and supervised the work. PR, QY, and SC prepared the manuscript with inputs from all authors.

## Supplementary Information

**Supplementary Table: Reagent and resource information.**

**Supplementary Data (provided in a zip file)**

**Supplementary Data 1: Metabolic gene candidate multi-tier selection information.**

**Supplementary Data 2: CRISPR-RNP gene editing indel detection and quantifications.**

**Supplementary Data 3: RNA-seq of *ADA* and *PDK1* KO and OE mutant T cells.**

**Supplementary Data 4: Metabolomics analyses in *ADA* and *PDK1* KO and OE mutant T cells.**

**Supplementary Data 5: Cytotoxicity assay data in *ADA* and *PDK1* KO and OE mutant CAR-T cells.**

**Supplementary Data 6: RNA-seq of *ADA* and *PDK1* KO and OE mutant CAR-T cells.**

**Supplementary Data 7: Metabolomics analyses in *ADA* and *PDK1* KO and OE mutant CAR-T cells.**

**Supplementary Data 8: Flow cytometry statistical analyses.**

## Figure Notes

Metabolomics data were analyzed by unpaired t test; indel and flow cytometry data were analyzed with Welch’s unpaired t tests; and tumor growth experiments were analyzed by 2-way ANOVA tests with Sidak’s multiple-comparison test. Flow cytometry plots shown with representative example data for each treatment. All analyses are presented with two-tailed results; error bars represent mean +/-SD; adjusted p values are FDR-corrected, unless stated otherwise. **** p < 1e-4, *** p < 1e-3, ** p < 0.01, * p < 0.05.

---

**Supplementary Fig. 1.**
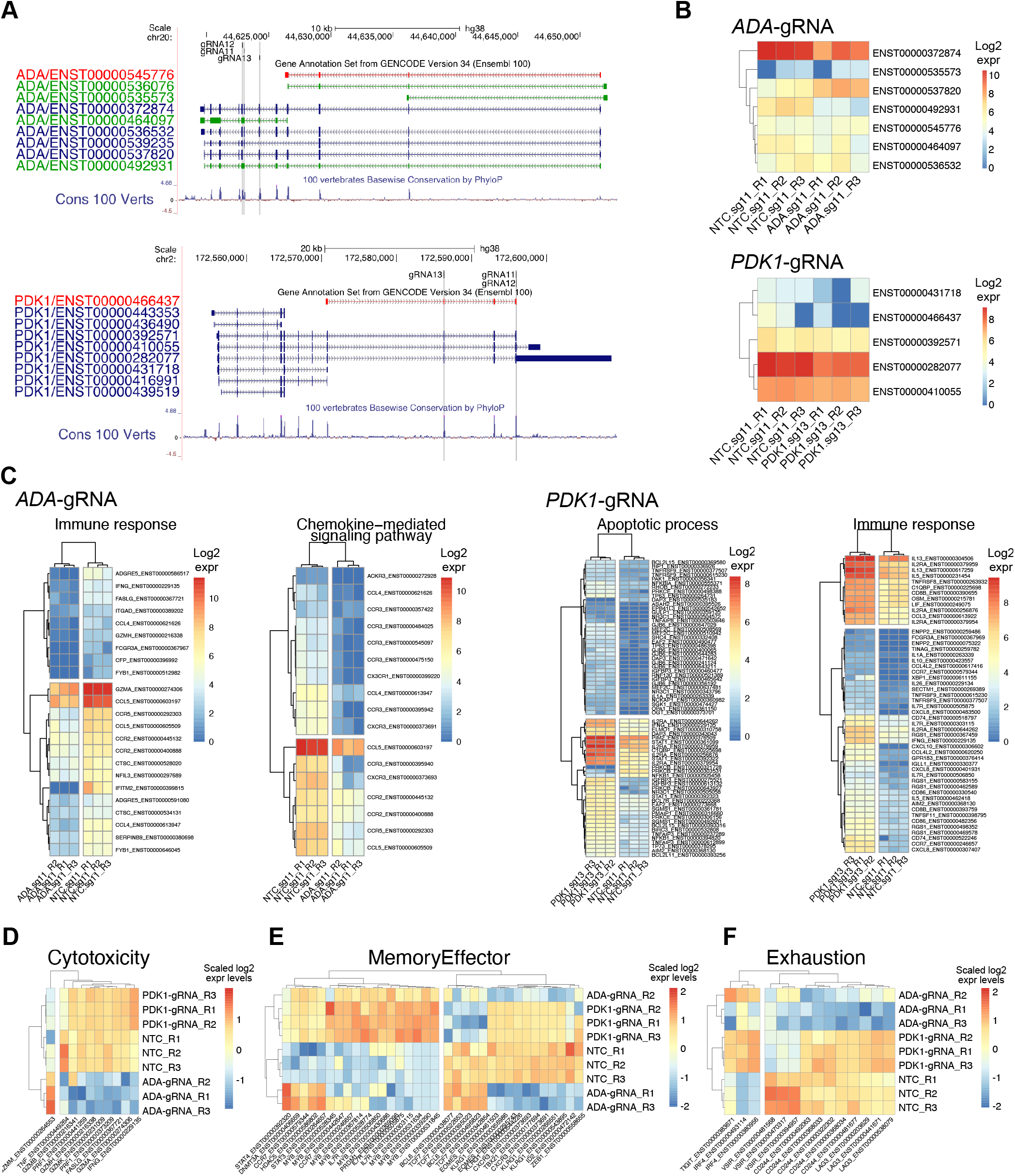

**Supplementary Fig. 2.**
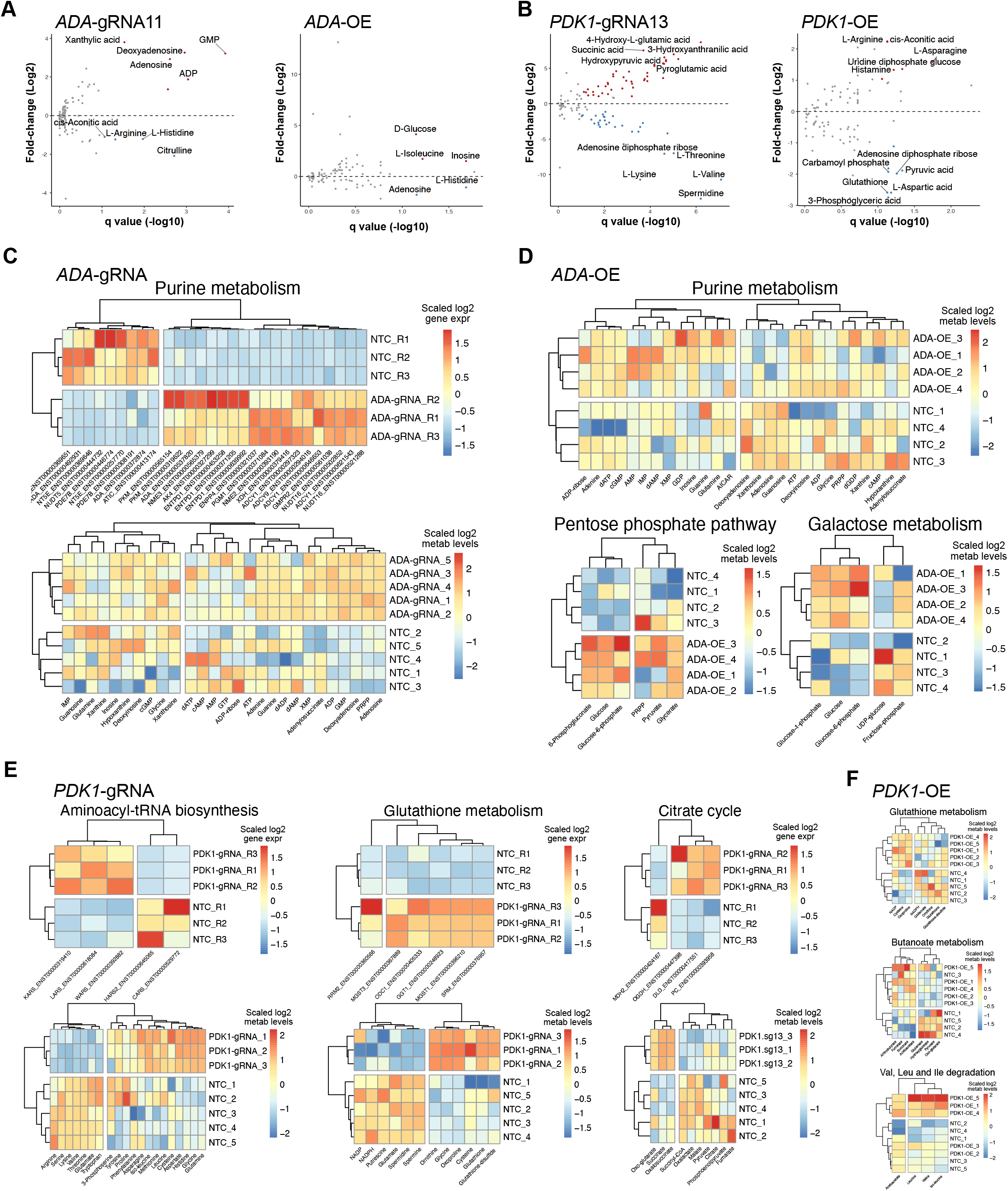

**Supplementary Fig. 3.**
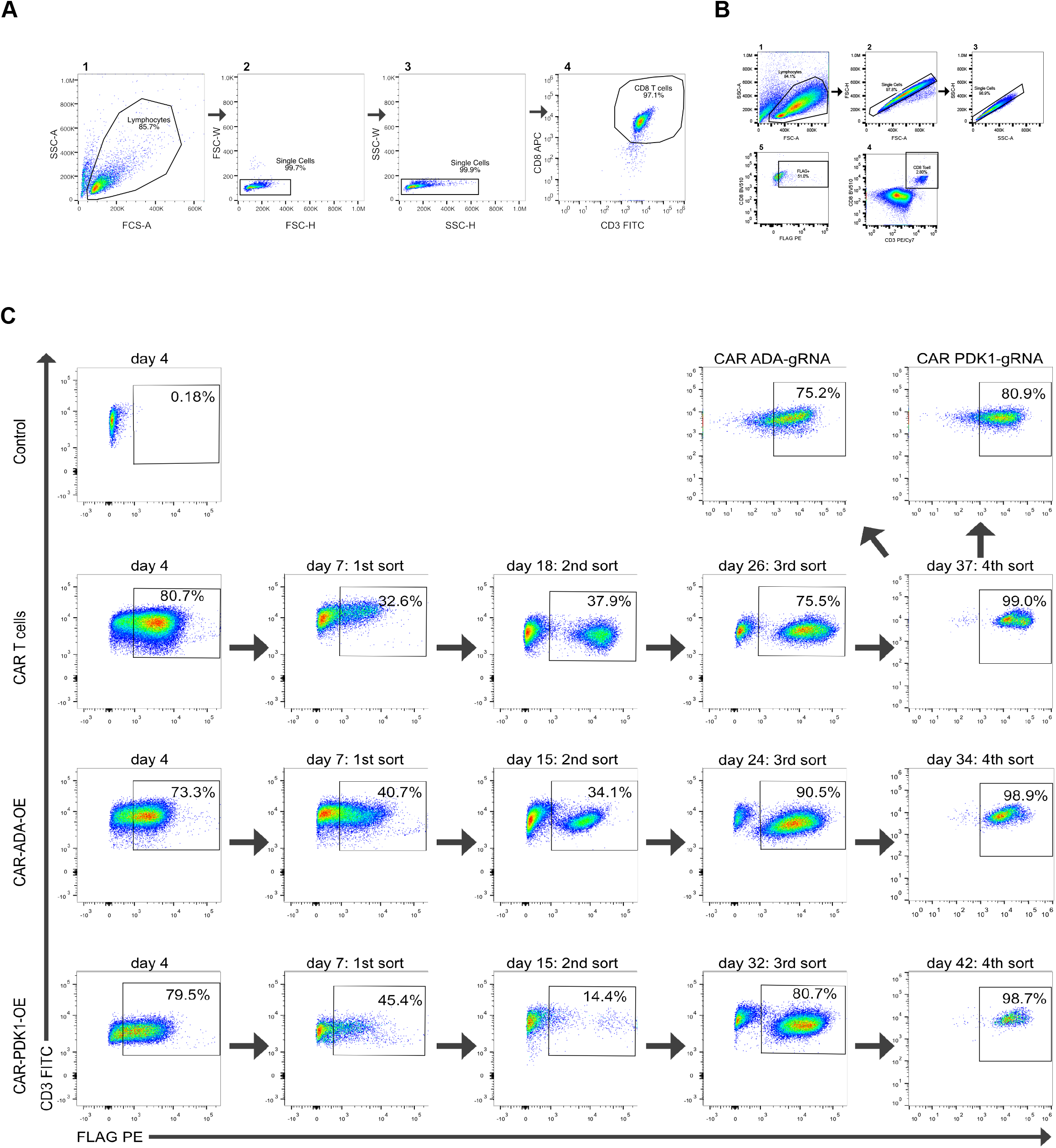

**Supplementary Fig. 4.**
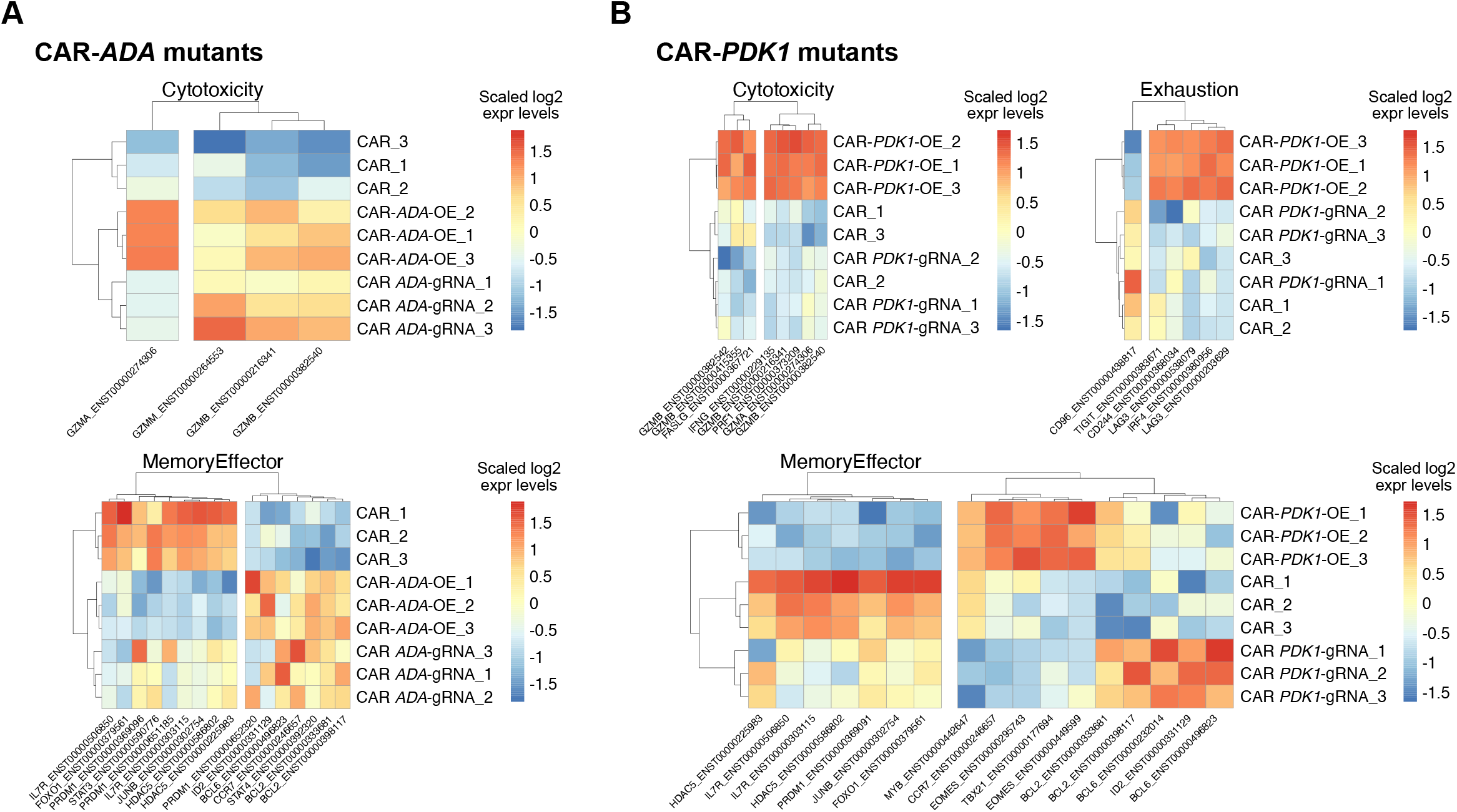

**Supplementary Fig. 5.**
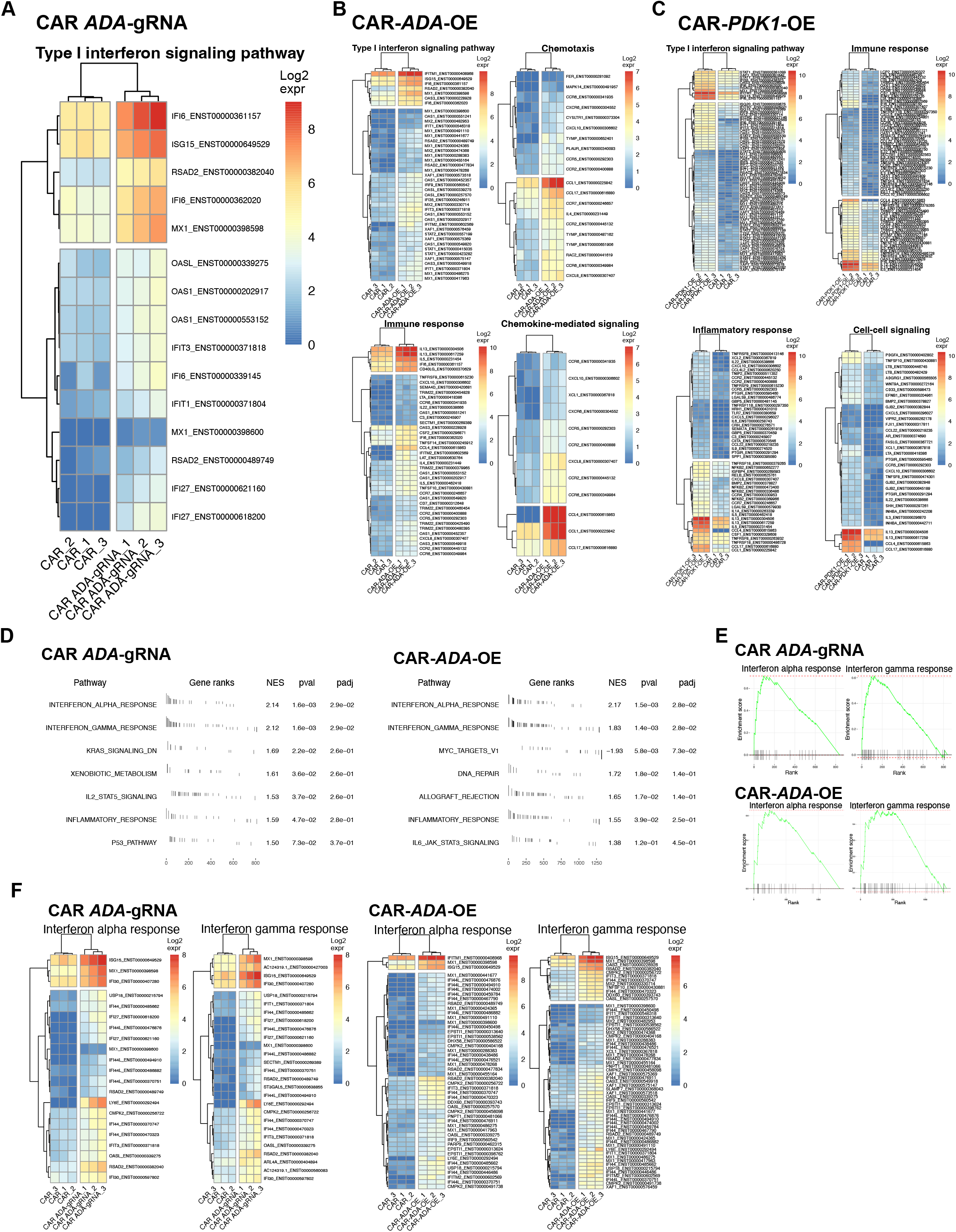

**Supplementary Fig. 6.**
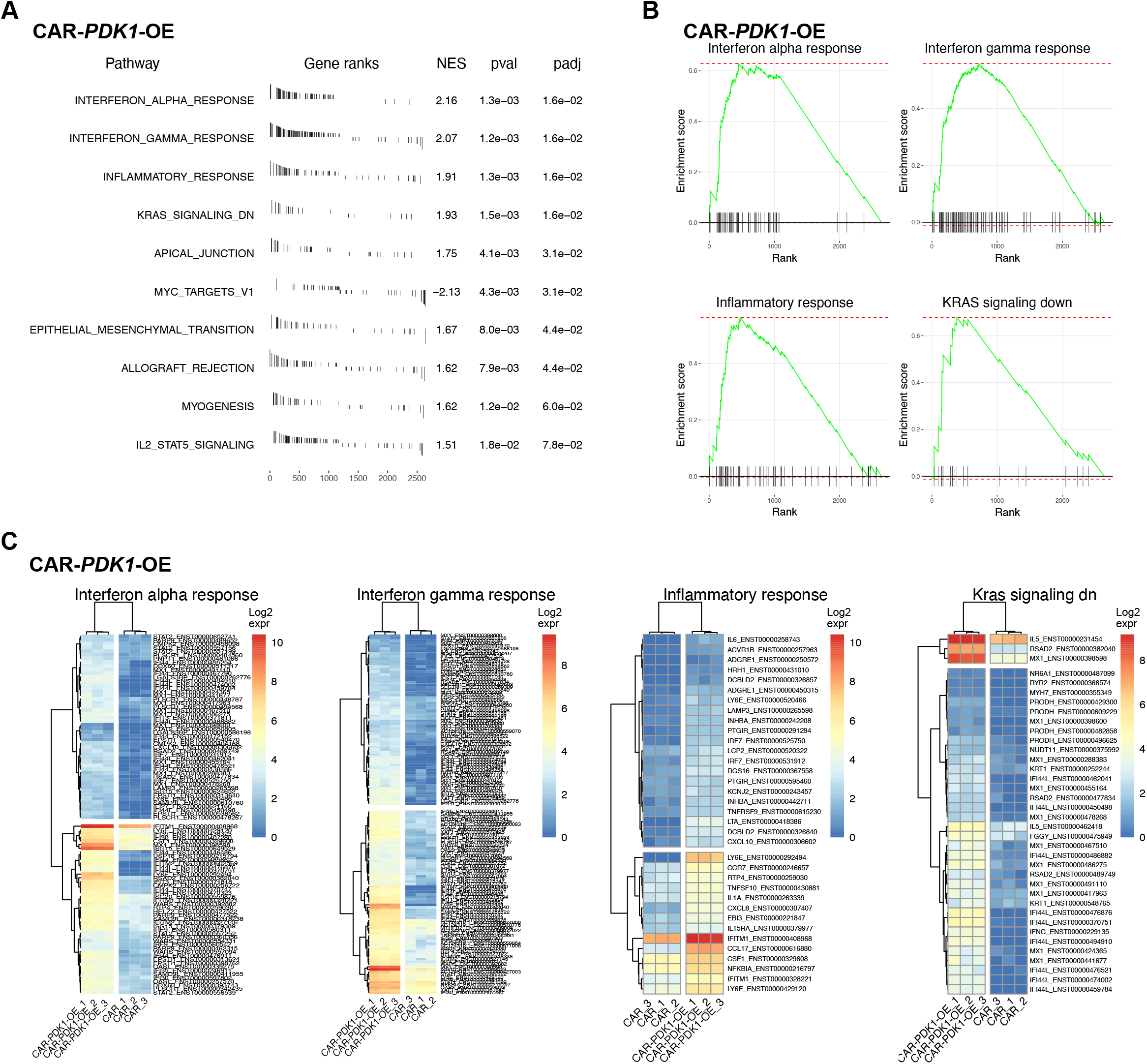

**Supplementary Fig. 7.**
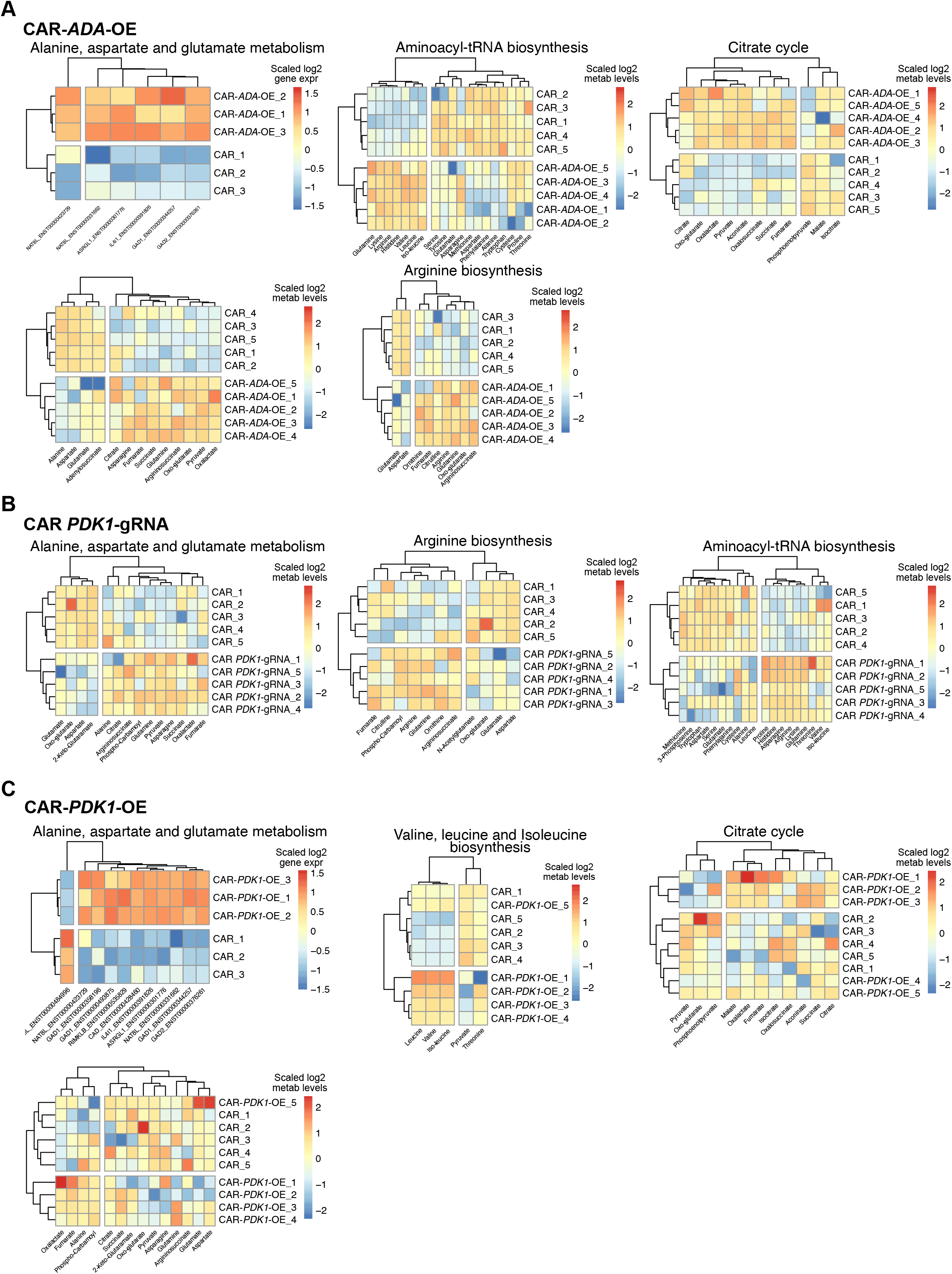

**Supplementary Fig. 8.**
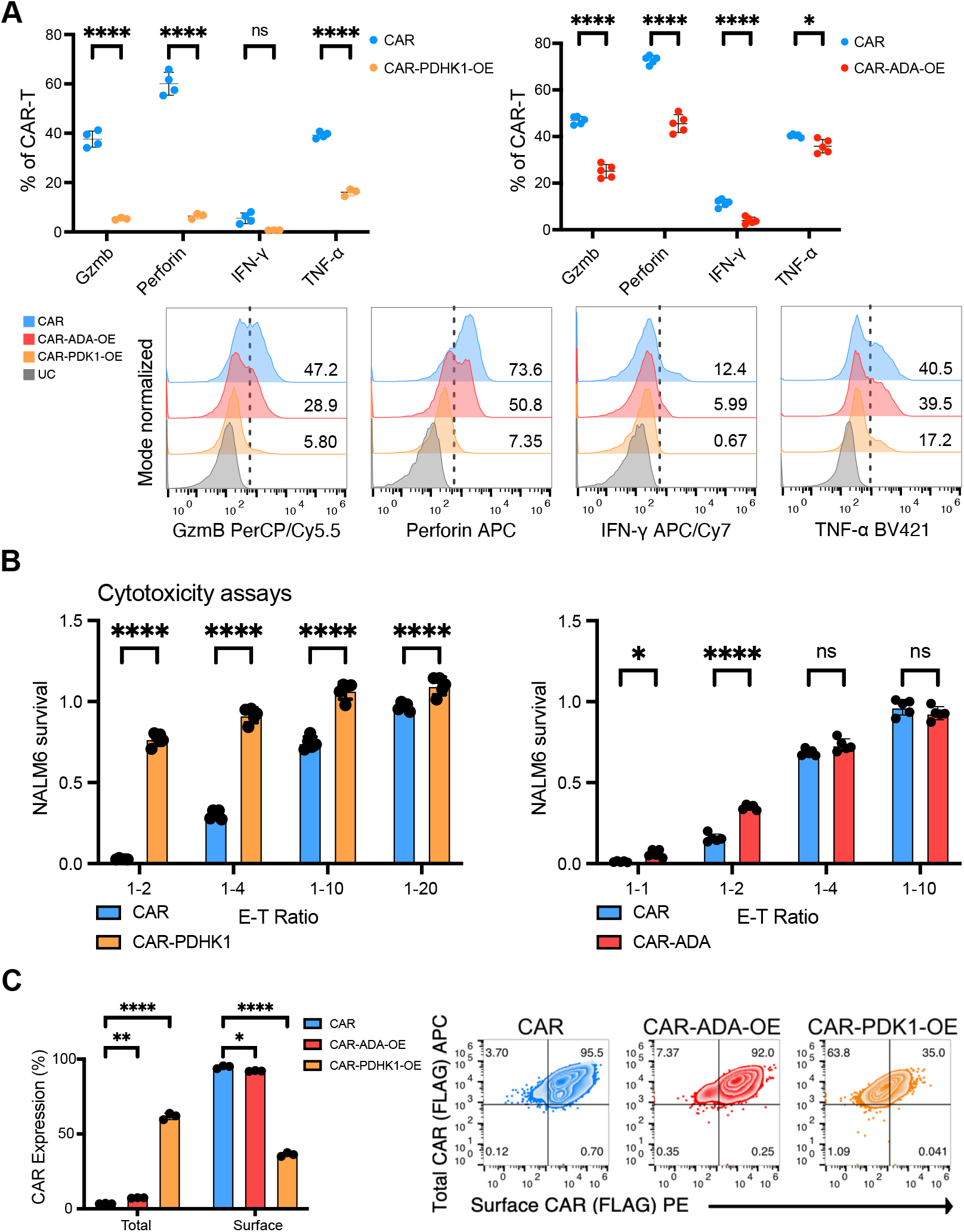

**Supplementary Fig. 9.**
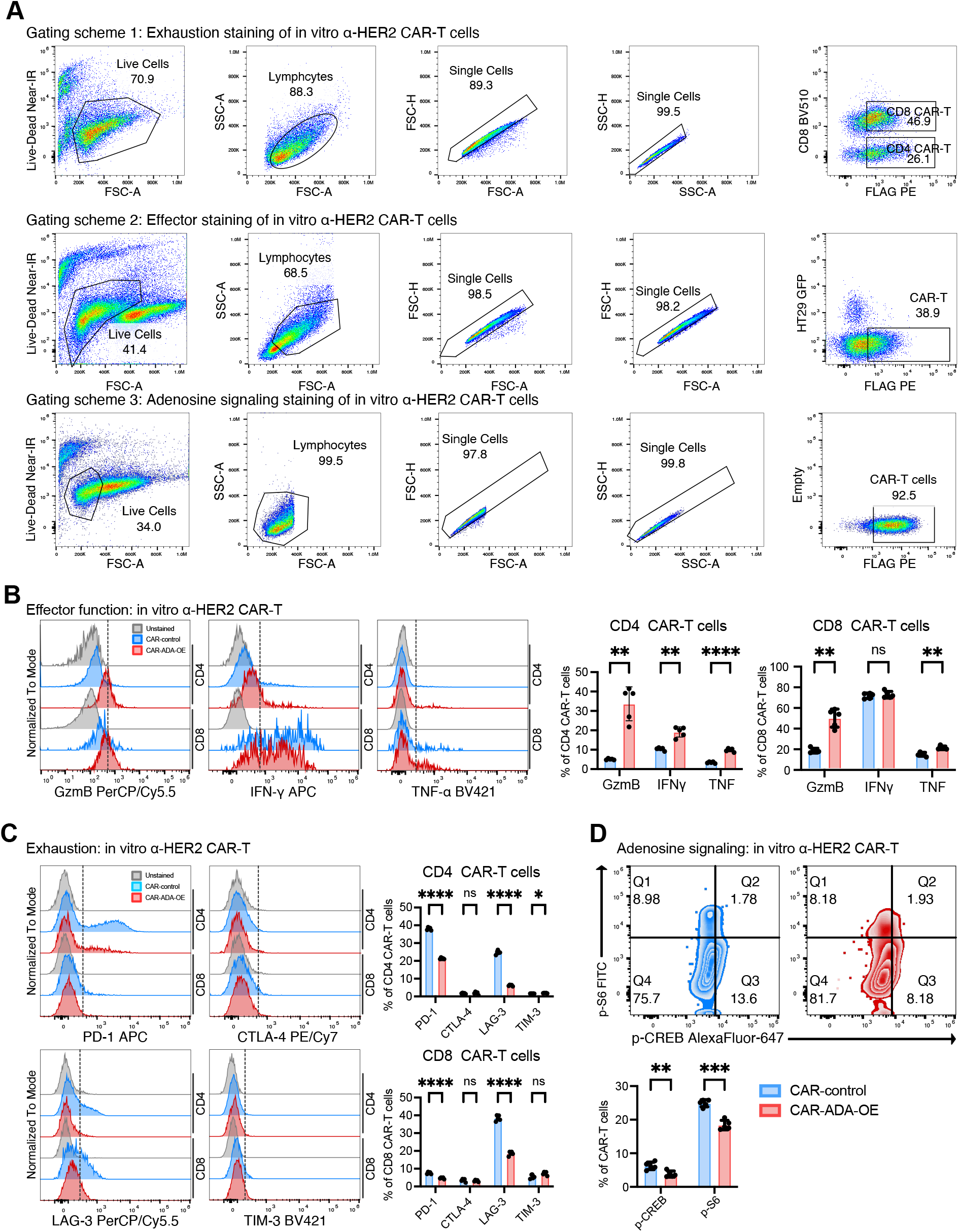

**Supplementary Fig. 10.**
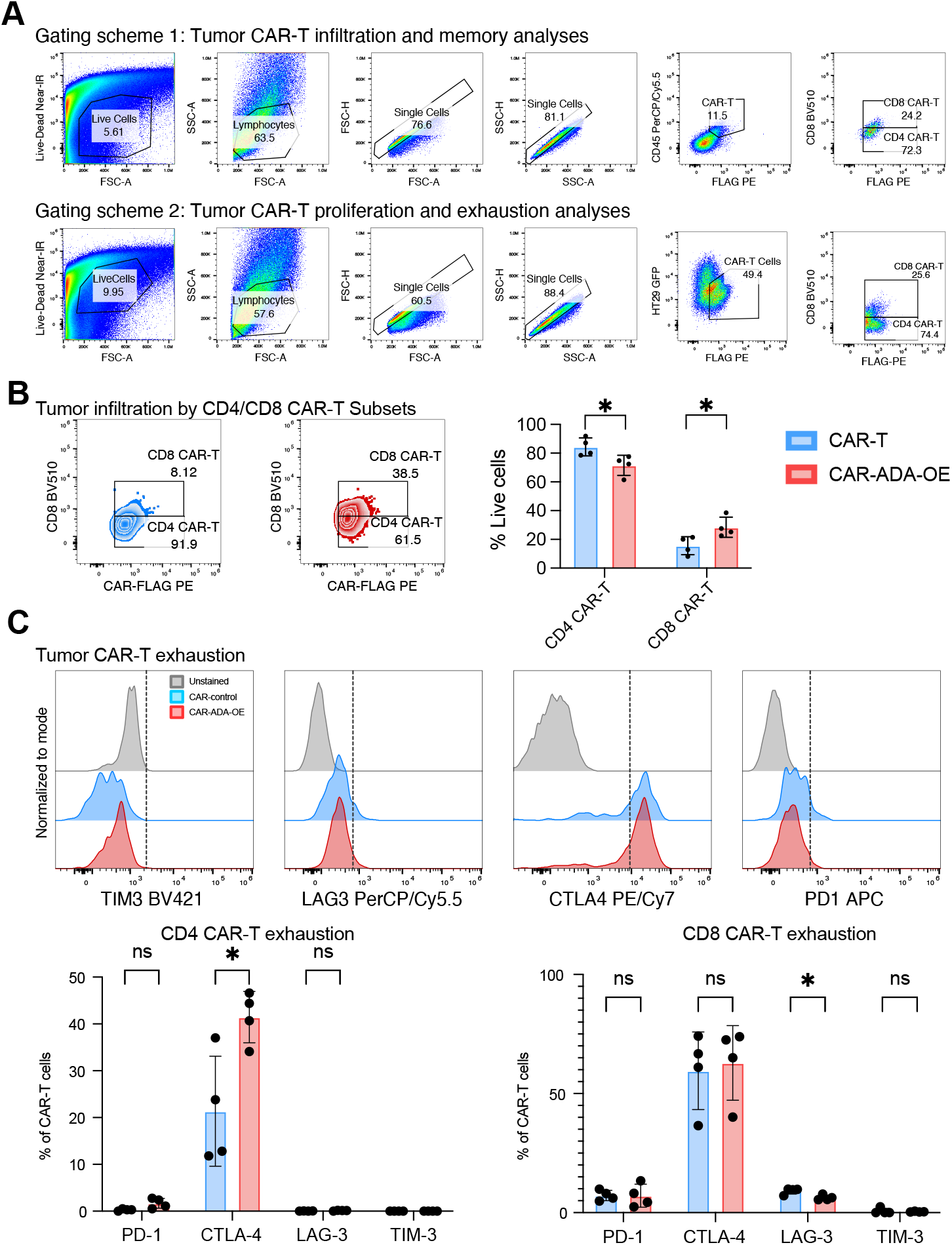

## Notes

### Competing Interest Statement

The authors have declared no competing interest.

